# The ataxia-telangiectasia disease protein ATM controls vesicular protein secretion via CHGA and microtubule dynamics via CRMP5

**DOI:** 10.1101/2024.06.26.600760

**Authors:** Marina Reichlmeir, Ruth Pia Duecker, Hanna Röhrich, Jana Key, Ralf Schubert, Kathryn Abell, Anthony P. Possemato, Matthew P. Stokes, Georg Auburger

**Affiliations:** Goethe University Frankfurt, University Hospital, Clinic of Neurology, Exp. Neurology, Heinrich Hoffmann Str. 7, 60590 Frankfurt am Main, Germany; Division for Allergy, Pneumatology and Cystic Fibrosis, Department for Children and Adolescence, Goethe-University, Frankfurt am Main, Germany; Institute for Experimental Pediatric Hematology and Oncology, Medical Faculty, Goethe-University Frankfurt, Komturstrasse 3a, 60528, Frankfurt am Main, Germany; Cell Signaling Technology, Inc., Danvers, MA 01923, USA

**Keywords:** PhosphoScan, Immobilized metal affinity chromatography, label-free mass spectrometry, co-immunoprecipitation, sodium arsenite, chloroquine, taxol, differential ultracentrifugation

## Abstract

The autosomal recessive disease ataxia-telangiectasia (A-T) presents with cerebellar degeneration, immunodeficiency, radiosensitivity, capillary dilatations, and pulmonary infections. Most symptoms outside the nervous system can be explained by failures of the disease protein ATM as Ser/Thr-kinase to coordinate DNA damage repair. However, ATM in adult neurons has cytoplasmic localization and vesicle association, where its roles remain unclear. Here, we defined novel ATM protein targets in human neuroblastoma cells and filtered initial pathogenesis events in ATM-null mouse cerebellum. Profiles of global proteome and phosphorylome - both direct ATM/ATR-phosphopeptides and overall phosphorylation changes - confirmed previous findings on NBN, MRE11, MDC1, CHEK1, EIF4EBP1, AP3B2, PPP2R5C, SYN1 and SLC2A1. Even stronger downregulation of ATM/ATR-phosphopeptides after ATM-depletion was documented for CHGA, EXPH5, NBEAL2 and CHMP6 as key factors of protein secretion and endosome dynamics, as well as for CRMP5, DISP2, PHACTR1, PLXNC1, INA and TPX2 as neurite extension factors. Prominent affection of semaphorin-CRMP5-microtubule signals and ATM association with CRMP5 were validated. As a functional consequence, microtubules were stabilized, and neurite retraction ensued. The ATM impact on secretory granules confirms previous ATM-null cerebellar transcriptome findings. Our study provides the first link of A-T neural atrophy to growth cone collapse and aberrant microtubule dynamics.

## 3. Introduction

The protein Ataxia Telangiectasia Mutated (ATM) belongs to the family of phosphoinositide 3-kinase (PI3K) related kinases (PIKKs, also including ATR and mTOR kinases), which are implicated in DNA damage response (DDR) and checkpoint signaling (Phan and Rezaeian, 2021; Savitsky et al., 1995a; Savitsky et al., 1995b). Disruption of the *ATM* gene and loss-of-function of ATM protein usually leads to the development of autosomal recessively inherited ataxia telangiectasia (A-T), a progressive disease presenting with early onset cerebellar degeneration, ocular telangiectasia, immunodeficiency with frequent pulmonary infections, and radiosensitivity leading to malignancies (Boder and Sedgwick, 1958; McKinnon, 2004).

The objective diagnosis of A-T is established by demonstrating increased blood levels of the fetal osmotic regulator AFP, which contrast with low levels of the adult osmotic regulator albumin and of total proteins (Ehlayel et al., 2014; Schieving et al., 2014; Szczawińska-Popłonyk et al., 2020). Many patients have short stature with lower fat-free mass and dyslipidemia, later developing insulin resistance with various endocrine deficits that are reflected by decreased blood levels of IGF-1 (insulin-like growth factor 1), GH (growth hormone), ACTH (adreno-corticotropic hormone) and vitamin D (Cirillo et al., 2018; Ehlayel et al., 2014; Kieslich et al., 2010; Kovacs et al., 1997; Nissenkorn et al., 2016; Petley et al., 2022; Pommerening et al., 2015; Ross et al., 2015; Rubio Pérez et al., 1978; Schalch et al., 1970; Schubert et al., 2009; Voss et al., 2014; Woelke et al., 2017). The conversely increased levels of TSH (thyroid stimulating hormone), FSH (follicle stimulating hormone) and LH (luteinizing hormone), which are all released from the pituitary gland, may be accompanied by deficient release of TRH (thyrotropin releasing hormone) and GnRH1 (gonadotropin releasing hormone 1, also known as LH-RH or FSH-RH) from hypothalamic neuroendocrine cells (Ammann et al., 1970; Zadik et al., 1978). Autopsy of A-T patient brains has shown the adenohypophysis cells to display cytomegaly with giant nuclei and vacuolization of the cytoplasm, whereas the hypothalamic dorso-medial and ventro-medial nuclei contained some multinucleated neurons that had the rough endoplasmic reticulum (also known as Nissl substance) relocalized to the cell periphery (BOWDEN et al., 1963; Kovacs et al., 1997). Ballooning degeneration of the spinal ganglia neurons, and cytomegaly with giant hyperchromatic nuclei were independently observed in adenohypophysis, thyroid, adrenal cortex and medulla, kidney, spleen, liver, dorsal root ganglia satellite cells, Schwann cells, capsular cells of myenteric plexus, epididymis, smooth muscle, heart, lung alveolar cells, and connective tissue (AGUILAR et al., 1968).

Most A-T features (e.g. immunodeficiency and radiosensitivity) can be explained by the well-known functions of ATM during the DDR. Particularly at DNA double-strand breaks (DSB) induced by ionizing irradiation (Suzuki et al., 1999), which are recognized by binding of the protein RAD50, the nuclease MRE11, and NBN as an activator of ATM, its downstream phosphorylation signals are crucial for the coordination of repair. After NBN-dependent autophosphorylation of ATM at serine (Ser) 1981 (You et al., 2005), the active ATM monomer, as a central component of the DDR, activates MDC1 to sustain the interactions (Lukas et al., 2004) and signals to various targets of the DDR pathway via its kinase function (Bakkenist and Kastan, 2003; Berkovich et al., 2007). While DDR is ongoing, the cell cycle will be halted by MDC1 and downstream CHEK1 signaling (Lou et al., 2003; Wang et al., 2011). ATM also modulates the nuclear versus cytoplasmic redistribution of protein phosphatase 2A (PP2A), which influences the autophosphorylation of ATM (Goodarzi et al., 2004) and coordinates mitosis versus cell growth (Sule et al., 2022). DSBs are then corrected mostly via homology-directed repair (HDR) (Nimonkar et al., 2011; Wawrousek et al., 2010) but also via non-homologous end joining (NHEJ) (Bennardo and Stark, 2010; IMAMICHI et al., 2014).

Given that the neurological features of A-T are usually one of the first signs for the disease seen from toddler stage on, manifesting in ataxia and indicating cerebellar dysfunction (Hoche et al., 2012), the reasons for the selective vulnerability of cerebellar neurons remain of outstanding interest. By the age of 10 years, abnormal movements get progressively worse, and patients usually become wheelchair-bound. Structurally, cerebellar dysfunction in A-T is due to atrophy mainly of granule neuron and Purkinje cell (PC) axons and dendrites, but also of dentate neurons and inferior olivary neurons (AGUILAR et al., 1968; Amirifar et al., 2019). However, the degeneration of post-mitotic cerebellar neurons, as well as the pathology of post-mitotic hypothalamic cells or adrenal medulla chromaffin cells that results in neuroendocrine deficits (Bunn et al., 2012; Constantin, 2011), cannot be explained by ATM functions in DDR. It has been demonstrated that ATM is mostly localized to the cytoplasm in adult cerebellar neurons (Oka and Takashima, 1998). Cytoplasmic localization of ATM was also demonstrated in the neuroblastoma cell line SH-SY5Y which shows chromaffin features (Hedborg et al., 2003; Wassberg et al., 1999), where translocation of ATM from the nucleus towards the cytoplasm was captured during the differentiation process (Boehrs et al., 2007).

Numerous studies have sought to elucidate the role of cytoplasmic ATM over the past decades. ATM was demonstrated to impact the trophic control over translation initiation via EIF4EBP1 (Mallory and Petes, 2000; Yang and Kastan, 2000), and to impact glucose transporter degradation (gene symbol SLC2A4 or SLC2A1) via the regulation of lysosomal number (Barlow et al., 2000; Cheng et al., 2020). ATM also localizes to endosomes, given that both *in vitro* and *in vivo* co-localization and co-immunoprecipitation studies revealed ATM association with β-adaptin (AP3B2) (Lim et al., 1998). ATM influence on the excitatory/inhibitory balance of neurons (Cheng et al., 2018) may be explained by its association with synaptic vesicles in co-localization with synapsin-1 (SYN1) and synaptobrevin-2 (VAMP2) (Li et al., 2009). Potential further cytoplasmic regulatory roles of ATM include the phosphorylation of cortactin (CTTN) as actin-binding protein to promote breast cancer cell migration (Lang et al., 2018). The actin-binding protein Drebrin (DBN) was also reported to be phosphorylated by ATM, as a pathway to protect synapses from oxidative stress (Kreis et al., 2019; Kreis et al., 2013). Actin cytoskeletal abnormalities are frequently found in several neurodegenerative diseases like Friedreich’s ataxia, Alzheimer’s Disease, and Huntington’s Disease (Bayot et al., 2013; Heredia et al., 2006; Wennagel et al., 2022). As in other neurodegenerative diseases, the extracellular appearance of the axonal cytoskeleton component neurofilament light chain (NFL) in patient blood serum was shown to reflect the severity and progression of A-T (Donath et al., 2021).

Efforts to model the neurodegenerative process of A-T in mice were hampered by the fact that ATM-kinase-dead mutants show embryonic lethality (Putti et al., 2021), while ATM-null mice show either no neurological phenotype until old age, or die from their immunological deficit after developing a thymoma at the age of 3-5 months, with only subtle dysfunction of the nervous system (Barlow et al., 1996; Chiesa et al., 2000; Lavin, 2013). This is in curious contrast to A-T patients where the phenotype is more severe in the absence of ATM, while patients with complete ATM protein that retains only residual kinase activity experience a longer survival and normal endocrinology (Verhagen et al., 2012). The lifespan of ATM-null mice was successfully extended to maximally 12 months, after substituting ATM deficient bone-marrow derived cells by ATM-competent cells via transplantation, making it possible to study the initial stage of cerebellar pathology by imaging techniques and global transcriptome profiling (Bagley et al., 2004; Canet-Pons et al., 2018; Duecker et al., 2021; Duecker et al., 2019; Pietzner et al., 2016; Pietzner et al., 2013; Reichlmeir et al., 2023), but a proteome and phosphoproteome profiling effort is lacking so far. The global transcriptome profiles of cerebella from ATM-null mice suggested abnormal signaling via synaptic vesicles, neuropeptides, and some neurotrophins (Reichlmeir et al., 2023).

Attempting to model the cerebellar atrophy of A-T *in vitro*, induced pluripotent stem cells, neural and olfactory stem cells were generated (Carlessi et al., 2013; Nayler et al., 2012; Nurieva et al., 2023; Ovchinnikov et al., 2020; Stewart et al., 2013). First attempts were made to achieve a cerebellar differentiation, however, their differentiation towards an authentic cerebellar fate as opposed to a more unspecific hindbrain fate is still a major challenge (Erceg et al., 2010; Lee et al., 2013; Nayler et al., 2017; Tao et al., 2010; Tzur-Gilat et al., 2013). Additionally, the biomaterial generated is usually insufficient for proteome/phosphoproteome profiling, so the characterization of these stem cell models remained focused on morphological or survival phenotypes, nuclear ATM function, or transcriptome profiles (Nayler et al., 2017; Stewart et al., 2013).

Here, we employed the human SH-SY5Y neuroblastoma cell line, which has features of chromaffin differentiation, to generate a stable knockdown of ATM (ATM-kd cells, shATM) by using shRNA. We characterized their molecular adaptation in the absence of stress, versus treatment with osmotic or oxidative stress, at the level of (i) global proteomics, (ii) direct ATM/ATR-phosphorylation targets, and (iii) overall differential phosphorylation profiling. This approach identified a high number of novel and consistently ATM-dependent factors together with confirmatory findings for most known ATM targets. In addition, ATM-null mice received bone-marrow transplantation to extend their survival until the stage of incipient cerebellar atrophy, aiming to define the earliest events in the molecular cascade of pathogenesis. Pathway enrichment statistics in these three-layer proteomic-phosphoproteomic profiles demonstrate that ATM-kd is accompanied by the enrichment of hypophosphorylations in proteins involved in microtubule dynamics and axonal differentiation, which is highly consistent with *in vivo* conditions. We validated ATM-dependent reduced protein levels and deficient secretion of the secretory granule biogenesis factor chromogranin A (CHGA), as a likely cause for the neuroendocrine deficits in A-T. Furthermore, we identified the collapsin response mediator protein 5 (CRMP5) in neural growth cones as ATM-dependent, as demonstrated by reduced phosphorylation at the typical ATM target sequence around phospho-Ser-538, accompanied by general protein reduction and by the stable association of ATM with CRMP5 in co-immunoprecipitation experiments. As phenotypic consequences, ATM-kd neuroblastoma cells displayed reduced neurite length and increased microtubule stability. This provides first evidence that impaired neuropeptide and neurotrophin secretion, neurite extension and synaptic pruning, as well as altered microtubule dynamics contribute to the selective vulnerability of neuroendocrine cells and cerebellar neurons at advanced stages of ataxia-telangiectasia.

## 4. Materials and Methods

### 4.1. Animal model of Ataxia-Telangiectasia

ATM-null mice were previously described (Barlow et al., 1996; Reichlmeir et al., 2023). Bone marrow transplantations were carried out as reported before (Reichlmeir et al., 2023).

### 4.2. Cell Culture

The stable *ATM* knockdown (kd) SH-SY5Y cell line (parental cell line ATCC: CRL-2266) was already published (Reichlmeir et al., 2023). In brief, stable ATM-kd cells were generated by lentiviral transduction of shRNA targeting *ATM* (hereafter referred to as shATM) compared to a control cell line targeting no known mammalian genes (non-target control, or NT CTRL). Cells were cultured in high glucose DMEM (Thermo Fisher Scientific, Waltham, Massachusetts, USA, 21969-035) supplemented with 10% fetal calf serum (FCS, Thermo Fisher Scientific, A3160802), 1% L-glutamine (Thermo Fisher Scientific, 25030-024), 0.1% penicillin / streptomycin (Thermo Fisher Scientific, 15140-122) and 1.25 µg/mL puromycin (Santa Cruz Biotechnology, Dallas, TX, USA, sc-108071). The cells were cultured at 37 °C and 5% CO_2_ in a humidified environment.

Chloroquine (CQ, Sigma-Aldrich, St. Louis, Missouri, USA, C6628) was administered to generate osmotic stress *in vitro*, which is known to activate ATM (Marceau et al., 2012; Qian et al., 2018). CQ was administered at a concentration of 20 µM for 24 hours with sterile water as control. Oxidative stress was generated *in vitro* by administration of 0.5 mM sodium arsenite (NaARS, Sigma-Aldrich, C6628) for 45 minutes with sterile water as control. Taxol (Cytoskeleton, Inc., Denver, Colorado, USA, TXD01) was used as a microtubule stressor. Taxol is an antitumor agent, causing microtubule assembly and stabilization (Schiff et al., 1979; Wani et al., 1971). Taxol was used at a concentration of 1 µM for 1 hour with dimethyl sulfoxide (DMSO, Sigma-Aldrich) applied as control.

Cells were harvested for protein or RNA analysis in PBS using cell scrapers. After centrifugation, pellets were frozen until usage for either reverse transcriptase real-time quantitative polymerase chain reaction (RT-qPCR), immunoblotting or co-immunoprecipitation (Co-IP).

### 4.3. (Phospho)-proteomics

Transient “kiss-and-run” interactions of the ATM kinase, mediating phosphorylations on target proteins at the S/T-Q motif were identified by ATM/ATR-motif specific phosphoproteome surveys. General phosphorylation differences were documented by global phosphoproteome profiles via IMAC technology.

For this purpose, the human neuroblastoma cells were first seeded at a density of 12 million cells in T125 flasks and allowed to settle overnight. Next, the cells were treated with CQ or control conditions as described above. 45 minutes before the end of the incubation time, the remaining cells were treated with NaARS as described above. Finally, the cells were harvested by trypsinization, pelleted by centrifugation, and washed once in PBS. After pelleting again, the cells were snap-frozen until mass spectrometry analysis.

For the *in vivo* analysis of phosphorylation events, aged (8-month-old) ATM-null mice (KO) and age- and sex-matched wildtype (WT) littermates (4 vs. 4) were sacrificed by decapitation, cerebella were removed and snap frozen in liquid nitrogen until mass spectrometry analysis.

The human cell lines and mouse tissues were prepared in urea lysis buffer (9 M urea, 20 mM HEPES pH 8.0 + phosphatase inhibitor cocktail) and shipped to Cell Signaling Technology (CST) for analysis. Cells and tissues were sonicated and centrifuged to remove insoluble material. Protein content was determined by Bradford assay and equal protein quantities from all samples were used for analysis. Samples were reduced with DTT and alkylated with iodoacetamide, then digested with trypsin (ATM/ATR Substrate) or LysC + trypsin (Fe-IMAC, Total Proteome) (CST, trypsin #56296, LysC #84748), purified over C18 columns (Waters) and enriched using the PTMScan ATM/ATR Substrate Motif (s/tQ) Kit (CST, #12267) or Fe-IMAC beads (CST, #20432) as previously described (Stokes et al., 2015). Samples were desalted over C18 tips prior to LC-MS/MS analysis. For Fe-IMAC and total proteome profiling, samples were labelled with Tandem Mass Tag reagents (ThermoFisher, #A52045) as previously described (Possemato et al., 2021). For total proteome profiling, samples were bRP fractionated collecting 96 fractions and concatenating non-sequentially to 12.

LC-MS/MS analysis was performed using a Thermo Orbitrap Fusion™ Lumos™ Tribrid™ mass spectrometer as previously described (Possemato et al., 2021; Possemato et al., 2017; Stokes et al., 2015) with replicate injections of each sample. Briefly, peptides were separated using a 50 cm x 100 µM PicoFrit capillary column packed with C18 reversed-phase resin and eluted with a 90-minute (ATM/ATR) or 150-minute (Fe-IMAC, total proteome) linear gradient of acetonitrile in 0.125% formic acid delivered at 280 nl/min. Tandem mass spectra were collected in a data-dependent manner using a 3 sec cycle time MS/MS method, a dynamic repeat count of one, and a repeat duration of 30 sec. Real time recalibration of mass error was performed using lock mass (Olsen et al., 2005) with a singly charged polysiloxane ion m/z = 371.101237. For Fe-IMAC and total proteome TMT samples, an MS3 method was used to reduce ion interference and ratio compression. Full MS parameter settings are available upon request.

MS spectra were evaluated using Comet and the GFY-Core platform (Harvard University) (Eng et al., 2013; Huttlin et al., 2010; Villén et al., 2007). Searches were performed against the most recent update of the Uniprot *Homo sapiens* and *Mus musculus* databases with a mass accuracy of +/-20 ppm for precursor ions and 0.02 Da for product ions. Results were filtered to a 1% peptide-level FDR with mass accuracy +/-5ppm on precursor ions and presence of a phosphorylated residue for enriched samples. Total proteome results were further filtered to a 1% protein-level FDR. Site localization confidence was determined using AScore (Beausoleil et al., 2006). All quantitative results for the ATM/ATR enrichment were generated using Skyline (MacLean et al., 2010) to extract the integrated peak area of the corresponding peptide assignments. Accuracy of quantitative data was ensured by manual review in Skyline or in the ion chromatogram files. All Fe-IMAC and total proteome quantitative results were generated in GFY-Core using signal:noise values for each peptide and summing individual signal:noise values for all peptides for a given protein/site for TMT data. The total signal:noise across each channel was normalized by dividing the maximum signal:noise across the channels by the total signal:noise in each channel and multiplying each individual channel by the resulting number (so the offset is 1.0 for the max channel). For all analyses, ratios for each binary comparison were generated, converted to log2 scale, and normalized via median log2 offset. Normalized log2 ratios were converted back to fold changes. p values were calculated using a 2-tailed t-test.

### 4.4. *In silico* analysis of OMICS results

The obtained OMICS data were analysed for candidate targets *in silico*. Phosphoproteomic data resulting from the ATM/ATR-motif specific screen in SH-SY5Y neuroblastoma *in vitro* model system and in mouse cerebella were first sorted for high confidence targets by only considering detected peptides with a coverage (% CV) of at least 50% and a maximum abundance of at least 500,000. Conditions could be compared as follows: shATM vs. NT CTRL; shATM+CQ vs. NT CTRL+CQ; shATM+NaARS vs. NT CTRL+NaARS; NT CTRL+CQ vs. NT CTRL; NT CTRL+NaARS vs. NT CTRL; shATM+CQ vs. shATM; shATM+NaARS vs. shATM. Analysis was done in three biological replicates for each condition. From the resulting data, the highest changed unique proteins, when comparing unstressed shATM vs. NT CTRL (cut-off: fold change +/-2.0) were analysed for candidate targets *in silico* by using PANTHER gene classification system (https://pantherdb.org/, last accessed on May 5, 2024) and Cytoscape visualization tool combined with STRING Protein-Protein Interaction Network (https://string-db.org/, last accessed on May 20, 2024). Phosphoproteomic data resulting from the IMAC survey from the same samples were studied regarding the highest changed unique proteins when comparing shATM vs. NT CTRL (cut-off: fold change +/-2.0) by using PANTHER gene classification system (https://pantherdb.org/, last accessed on May 5, 2024). Protein-protein interactions and pathway enrichments were initially assessed and visualized via the STRING Heidelberg webpage (https://string-db.org/, last accessed on May 20, 2024), using default parameters, with the output being added to the original data contained in Supplementary tables as separate tabulators. Experimental observations of ATM/ATR phosphopeptide dysregulations were assessed if the amino acid sequence matches predictions as classical ATM targeting motif, employing the Cell Signaling Technology webtool (https://www.phosphosite.org/, last accessed on May 20, 2024).

*In vivo* phosphoproteomic data were analysed *in silico* by Cytoscape visualization tool combined with STRING Protein-Protein Interaction Network (https://pantherdb.org/, last accessed on May 5, 2024), including all significant hits in the analysis.

Volcano plots were generated using Microsoft Excel and Graphpad Prism (Version 8). Log_2_ fold change cutoffs were -1 and 1 for down- and up-regulations, respectively, and a –log_10_ (p-value) cut-off for significance at 1.3. For ATM/ATR and IMAC analyses, only the phosphorylation sites with the highest fold changes for each peptide were used in the visualizations.

### 4.5. RT-qPCR

Total RNA was isolated from cell pellets using TRI Reagent (Sigma-Aldrich), following the manufacturer’s protocol. Subsequent cDNA synthesis was performed using the SuperScript IV Kit (Invitrogen, Carlsbad, California, USA). For this, 1 µg RNA was first cleared from residual genomic DNA via incubation with ezDNase (Invitrogen). Then, reverse transcription was performed following the manufacturer’s instructions. Transcript levels were analysed by RT-qPCR using TaqMan Gene Expression Assays^TM^ (Thermo Fisher Scientific). This was done by using cDNA from 10 ng total RNA with 2x FastStart Universal Probe Master ROX (Roche, Basel, CHE) and the corresponding TaqMan Assay in a StepOnePlus Real-Time PCR Cycler (Applied Biosystems, Waltham, Massachusetts, USA). Data were assessed using the 2^-ΔΔCt^ method (Schmittgen and Livak, 2008). Data are displayed as fold changes to the unstressed NT CTRL condition. Human transcripts were quantified using the following TaqMan Assays: *ATM* – Hs01112311_m1; *DPYSL5* – Hs00184106_m1; *KIF1A* – Hs00987720_m1; *KIF21B* – Hs01118430_m1; *KIF26B* – Hs00215977_m1; *MAP2* – Hs00258900_m1; *SEMA3A* – Hs00173810_m1; *TUBB3* – Hs00801390_s1. *TBP* transcripts (Hs9999910_m1) served as a normalizer.

### 4.6. Immunoblotting

To assess protein levels in SH-SY5Y cells, samples were lysed for 30 minutes on ice in RIPA buffer (50 mM TRIS/HCl pH 8.0; 150 mM NaCl; 1% NP-40; 0.5% sodium deoxycholate; 0.1% sodium dodecylsulfate, SDS) supplemented with HALT phosphatase inhibitors (Thermo Fisher Scientific) and cOmplete proteinase inhibitors (Roche). Lysis was completed by brief sonication and removal of cell debris via centrifugation. Protein concentration was determined via Pierce BCA Protein Assay Kit (Thermo Fisher Scientific) following the manufacturer’s instructions. SDS-polyacrylamide gel electrophoresis (PAGE) was performed using 25 µg denatured (90 °C, 5 minutes) total protein, except stated otherwise. Transfer was done on 0.2 µm nitrocellulose membranes (Bio-Rad, Hercules, California, USA) and blocking was done in 5% bovine serum albumin (BSA, Carl Roth GmbH, Karlsruhe, GER) in TBS-buffer supplemented with 0.1% Tween-20 (TBS-T, Sigma-Aldrich) for one hour. Primary antibodies were incubated over night at 4 °C in blocking buffer. Primary antibodies were: anti-ATM (Cell Signalling Technology, Danvers, Massachusetts, USA, #2873); anti-CRMP5 (Proteintech, Chicago, Illinois, USA, 10525-1-AP); anti-CHGA (Cell Signalling Technology, #60893S); anti-TUBB3 (Invitrogen, MA1-118); anti-GAPDH (Sigma-Aldrich, CB1001); anti-STMN1 (Proteintech, 11157-1-AP); anti-phospho-STMN1-Ser16 (Cell Signalling Technology, #3353).

Subsequently, membranes were subjected to the respective fluorescent secondary antibody for one hour in TBS-T and finally imaged in LI-COR Odyssey Infrared Imager. Secondary antibodies were: IRDye 800CW goat anti-rabbit (LI-COR, Lincoln, Nebraska, USA, 926-32211); IRDye 680RD goat anti-rabbit (LI-COR, 926-68071); IRDye 800CW goat anti-mouse (LI-COR, USA, 926-32210); IRDye 680RD goat anti-mouse (LI-COR, 926-68070).

Quantification was done by densitometric analysis in ImageStudio Software using GAPDH as a normalizer. The protein content was quantified relative to the unstressed NT CTRL cells and is displayed as fold-change (FC).

### 4.7. Subcellular fractionation

To investigate ATM specifically in the cytoplasmic compartment of SH-SY5Y, fractionations were performed as previously described (Adrain et al., 2001; Reichlmeir et al., 2023). Protein concentration was determined by BCA assay (Thermo Fisher Scientific) following the manufacturer’s instructions. ATM protein interactions in the cytoplasmic compartment were analysed by subjecting cytoplasmic fractions to co-immunoprecipitations (Co-IP).

### 4.8. Immunofluorescence and Imaging

For immunofluorescence analysis, cells were first seeded at a density of 10,000 cells in 96-well black clear-bottom plates (Greiner BioOne GmbH, Frickenhausen, GER). The next day, cells were treated with Taxol as described above and finally fixed for 15 min in 4% PFA and washed in PBS. Cells were then permeabilized for 15 min in 0.1% Triton-X-100 / PBS. Following five washes with PBS, cells were blocked in antibody dilution buffer (ADB) (0.9% NaCl, 10 mM Tris/HCl pH 7.5, 5 mM EDTA, 1 mg/mL BSA) for 30 minutes. After that, the primary antibodies were incubated in ADB over night at 4 °C. Primary antibodies were anti-CRMP5 (Proteintech, 10525-1-AP) and anti-TUBB3 (Invitrogen, MA1-118).

After washing the cells again using PBS, the respective secondary antibodies were incubated in ADB at room temperature in the dark for 90 min together with 0.5 µg/mL 4′,6-diamidino-2-phenylindole (DAPI, Sigma Aldrich). Before imaging the cells were washed again using PBS. Secondary antibodies were AlexaFluor488 goat anti-rabbit (Invitrogen, A11034) and AlexaFluor594 goat anti-mouse (Invitrogen, A11032).

Imaging was conducted using the ImageXpress Micro XLS Widefield High-Content Analysis System (Molecular Devices, San Jose, California, USA), with 60x magnification. Per well, 64 sites were recorded, and images were subsequently processed with the MetaXpress® Software (Molecular Devices). Formatting of images was performed using ImageJ.

### 4.9. Quantification of fluorescence intensity

Fluorescence levels were quantified using ImageJ on ten respective images in three independent experiments per condition. For this, areas that contain cells in the green channel were first selected by using Auto Threshold function using the “Li” threshold filter algorithm. Then outlines of the detected cells were added as region of interest (ROI) by using the Analyze Particles Function. Particles were considered at a size of at least 50 pixels to exclude stained debris. Then, the mean grey value was measured in these areas in the green channel and in the red channel. The mean fluorescence intensity values of different ROIs in one picture were then calculated.

### 4.10. Co-immunoprecipitations

For co-immunoprecipitations, 800 - 1000 µg protein in cytoplasmic subcellular fractions of SH-SY5Y cells were incubated with either 10 µg anti-ATM antibody (Novus Biologicals, Littleton, Colorado, USA, NB100-220) or 10 µg of mouse IgG1 isotype control antibody (Cell Signalling Technology, #5415) in binding buffer consisting of CEB fractionation buffer supplemented with HALT phosphatase inhibitors (Thermo Fisher Scientific) and cOmplete proteinase inhibitors (Roche) over night at 4 °C with head to tail rotation. When using CRMP5 as prey protein, lysates were incubated with 8 µg anti-CRMP5 antibody (Proteintech, 10525-1-AP) or 8 µg of respective IgG isotype control antibody (Cell Signalling Technology, #2729). For the input control, 5% lysate was denatured at 90 °C for 5 min. The next day, the formed antibody-immunocomplexes were incubated with 100 µL Protein G magnetic Dynabeads^TM^ (Invitrogen, 10004D) for 1 h at room temperature with head-to-tail rotation allowing the complexes to bind the magnetic beads. Beads were then washed five times using PBS supplemented with HALT phosphatase inhibitors (Thermo Fisher Scientific) and cOmplete proteinase inhibitors (Roche). Finally, bound proteins were eluted from the magnetic beads by using 2x SDS sampling buffer. Eluted proteins were then denatured at 90 °C for 5 min and loaded in equal amount to two SDS-PAGE gels. Detection of the pulled proteins was performed by immunoblotting as described above. Detection Antibodies were anti-ATM (Cell Signalling Technology, #2873), anti-CRMP5 (Proteintech, 10525-1-AP) and anti-SPAST (Proteintech, 67361-1-Ig).

### 4.11. Quantification of microtubule content and free tubulin

The microtubule/tubulin *in vivo* assay kit (Cytoskeleton, Inc., #BK038) was performed to determine the amount of microtubule content as opposed to free tubulin *in vitro*. This was done by first seeding 1.5 million cells in 6 cm culture dishes. After 24 hours the cells were treated with CQ to elicit osmotic stress as described above. Additionally, the cells were also treated with 1 µM Taxol for 1 hour (provided in the kit) as a positive control for microtubule stabilization, while DMSO served as a control. After that, the cells were collected as suggested by the manufacturer. A first centrifugation step (1000 rcf, 5 min, 37 °C) yielded the low-speed pellet (LSP) fraction, which was resuspended in 2x SDS sample buffer at 1.2 volumes to the original amount of lysis buffer applied. The low-speed supernatant was then subjected to ultracentrifugation (100,000 rcf, 60 min, 37 °C) to separate microtubules to the pellets and unpolymerized tubulin to the supernatant. Finally, the high-speed supernatant (HSS) was removed and 5X SDS sampling buffer was added. Finally, the microtubules in the high-speed pellet (HSP) were depolymerized in depolymerization buffer, in an equal volume to the original lysis buffer applied. An appropriate amount of 5X SDS sampling buffer was added. An incubation was done for 15 minutes at room temperature with frequent pipetting to shear the microtubule pellet.

For quantification, the LSP, HSP and HSS fractions samples were loaded at a volume of 10 µL and separated by SDS-PAGE and detected as described above. Tubulin was detected by the included primary antibody. ATM (Cell Signalling Technology, #2873) and CRMP5 (Proteintech, 10525-1-AP) were also checked for their appearance in different fractions. Secondary antibodies were IRDye 800CW donkey anti-goat (LI-COR, 926-32214) and IRDye 680RD goat anti-rabbit (LI-COR, 926-68071). Quantification was done by densitometric analysis in ImageStudio Software. Protein content was quantified relative to the LSP of unstressed NT CTRL cells and displayed as fold-change.

### 4.12. Analysis of protein secretion

Secreted proteins were analysed in the cell culture supernatant. For this purpose, cells were seeded at a density of 4 million cells in 10 cm culture dishes and subsequently treated with CQ to elicit osmotic stress as described above. After the 24 h incubation period, the conditioned medium was harvested and centrifuged at 200 rcf for 3 minutes to remove cell debris. Then, exactly 9 mL of the supernatant was added to Amicon Ultra-15 Centrifugal Filter Devices (Merck Millipore, Burlington, Massachusetts, USA, 30K cut-off, UFC903008), which were equilibrated by centrifugation of 2 mL PBS (4000 rcf, 30 min, 4 °C) prior to use. The supernatant was finally concentrated by centrifugation (4000 rcf, 1 h, 4 °C). Exactly 12.5 µL of the concentrate was denatured (90 °C, 5 min) and subsequently separated by SDS-PAGE as described above. After the transfer, total protein was stained as a normalizer using the Revert 700 Total Protein Stain Kits (LI-COR, 926-11010) following the manufacturer’s instructions. Membranes were then blocked as described above and antibody staining was performed as for standard immunoblotting procedure (described above).

### 4.13. Quantification of neurite length

For the morphological analysis, cells were first seeded at an amount of 150,000 cells per well in 12-well plates. After 24 hours, the cells were treated with either 20 µM chloroquine (24 h) or 1 µM Taxol (1 h) as described above. Then, the cells were washed in PBS and stained with Hoechst-33258 (Invitrogen) for 10 minutes. After this, the cells were washed again in PBS and finally fixed by application of 4% PFA for 10 minutes. After three washes in PBS, the cells were imaged using the EVOS M5000 microscope (Thermo Fisher Scientific) with 20x objective. Pictures were taken in transmitted light channel and DAPI channel. For each condition, three wells were imaged as replicates, with five spots imaged for each condition. Measurement of neurite length was carried out using ImageJ software. From each picture, ten neurites were measured beginning from the apical part of the nucleus until the end of the process.

### 4.14. Statistics

Data were statistically analysed using GraphPad Prism 8 Software. Grouped data were assessed via 2-way ANOVA followed by Sidak’s post-hoc test for multiple comparisons. Comparisons of more than four groups in the quantification of tubulin content was performed with multiple unpaired t-tests without correction for multiple testing. Each fraction was studied individually without assuming consistent standard deviation. Two groups were analysed using unpaired t-test with Welch’s correction. Asterisks represent significance (* = p ≤ 0.05, ** = p ≤ 0.01, *** = p ≤ 0.001, **** = p ≤ 0.0001). P-values 0.05 < p < 0.10 were considered as a statistical trend (T). Data are displayed as mean ± standard error of the mean (SEM) without additional single values. Protein and transcript ratios are displayed as fold-changes, relative to the untreated control condition. Microtubule content quantification is displayed as fold-changes, relative to the untreated control condition, LSP fraction.

## 5. Results

### 5.1. ATM-kd neuroblastoma cell profiling with proteomics reveals novel ATM-dependent factors involved in secretory vesicles and tubulin-actin cytoskeleton regulation

As stated previously, ATM might serve as a platform for protein-protein interactions, in addition to functioning as kinase (Reichlmeir et al., 2023). Therefore, we performed global proteomics profiling *in vitro* in SH-SY5Y neuroblastoma cells with lentiviral transduction of either NT CTRL shRNA plasmid or shATM knockdown plasmid **(Fig. 1A, Supplementary Table S1)**. Knockdown efficiency and ATM cytoplasmic localization were confirmed previously (Reichlmeir et al., 2023). In addition, we applied the osmotic stressor CQ and the oxidative stressor NaARS to mimic aging stress *in vitro*. We performed *in silico* gene ontology enrichment analysis using PANTHER for downregulated and upregulated proteins with a fold change of at least +/- 2-fold when comparing only shATM vs. NT CTRL conditions **(Fig. 1B)**. For downregulated proteins we found enrichments e.g. for “regulation of neuron projection development (GO:0010975)” biological process, “mu-type opioid receptor binding (GO:0031852)” molecular function and “postsynaptic intermediate filament cytoskeleton (GO:0099160)” and “chromaffin granule membrane (GO:0042584)” cellular components. For upregulations however, we found enrichments for e.g. “protein targeting to lysosome involved in chaperone-mediated autophagy (GO:0061740)” biological process and “actin cap (GO:0030478)” cellular component. Depletion of ATM protein was confirmed in this global proteome screen. Beyond depletion of the known ATM interactors AP3B2, NBN and CHEK1, prominent downregulations were detected for CRMP5 (*DPYSL5* gene symbol) microtubule binding protein and CHGA secretory protein **(Fig. 1C)**. Subsequent validation efforts and analysis of extracellular CHGA in the cell culture supernatant demonstrated significant reduction of CHGA secretion from neuroblastoma cells upon loss of ATM **(Fig. 1D)**. These data demonstrate that downregulations are evident for proteins that regulate the tubulin-actin cytoskeleton and secretory granules.

**Figure 1:**
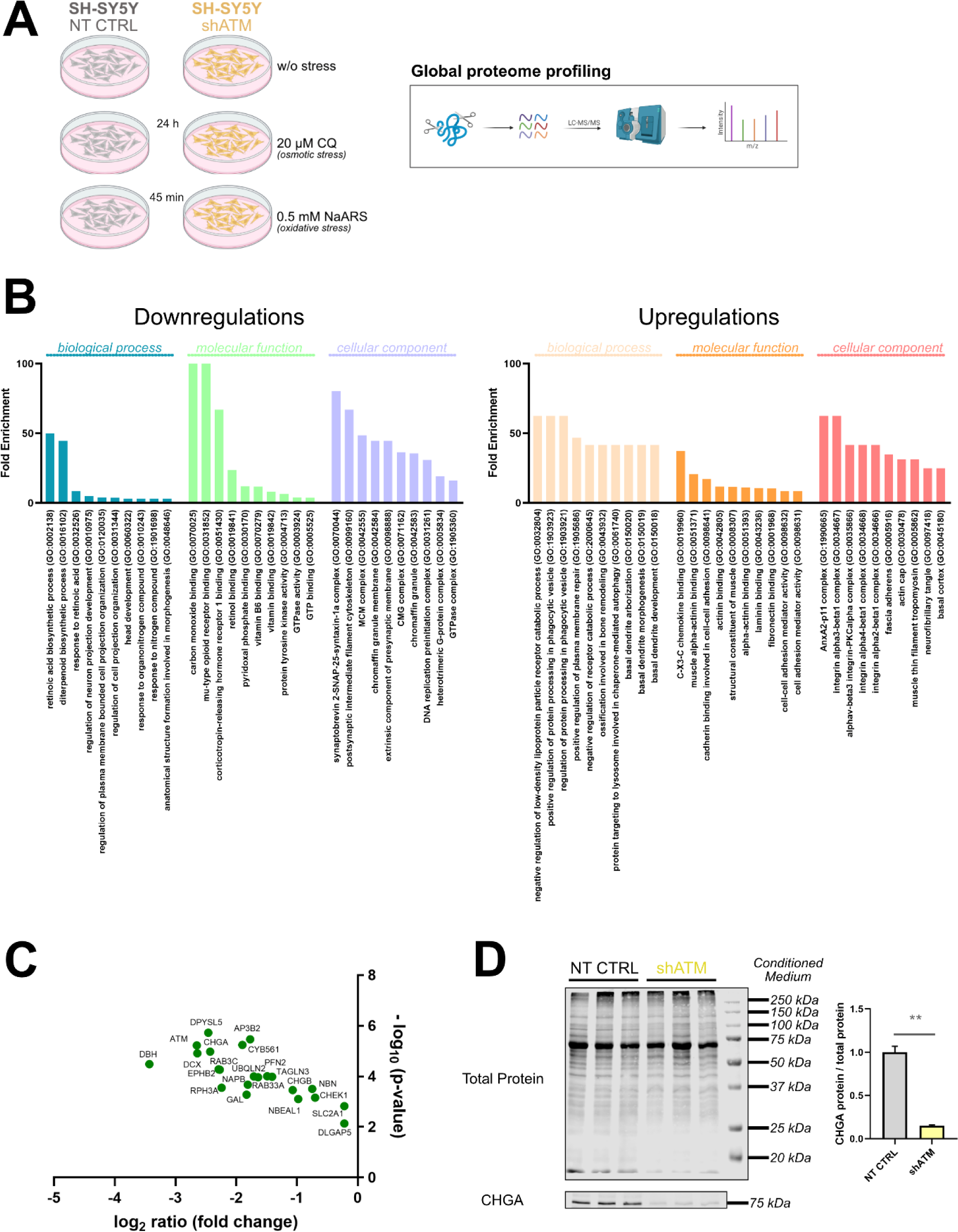
Total proteome profiling of SH-SY5Y neuroblastoma cells with ATM knockdown reveals secretory deficit upon ATM loss. **(A)** Workflow of the global proteome survey *in vitro*. SH-SY5Y NT CTRL cells and shATM cells (knockdown, ATM-kd) were subjected to quantitative detection of peptides by LC-MS/MS. Protein abundance was observed for (i) unstressed conditions, (ii) upon osmotic stress by administration of CQ, and (iii) upon oxidative stress by administration of NaARS. Each comparison represents three NT CTRL vs. three shATM biological replicates. **(B)** Functional enrichment of biological processes, molecular functions and cellular components for downregulations (left, green and blue shades) and upregulations (right, orange shades) with a fold change of at least ± 2.0 determined by PANTHER classification system (https://pantherdb.org/) when comparing NT CTRL and shATM conditions. **(C)** Volcano plot of selected downregulated proteins with fold change (calculated as log2 ratio) plotted against p-value (calculated as –log10). **(D)** Immunoblots for the quantification of secreted CHGA protein in cell culture medium with total protein staining as normalizer, and bar graphs with densitometric quantification of CHGA protein to total protein levels (n=3). Bar plots represent mean + SEM. Asterisks reflect significance: * = p ≤ 0.05, ** = p ≤ 0.01, *** = p ≤ 0.001, *** = p ≤ 0.0001; T = p ≤ 0.1, ns = non-significant. Abbreviations: CQ = chloroquine, NaARS = sodium arsenite, CHGA = chromogranin A.

### 5.2. ATM-kd neuroblastoma cell profiling with two phosphoscans reveals novel ATM kinase targets, affecting secretory vesicles and tubulin-actin cytoskeleton regulation

To identify novel cytoplasmic kinase targets of ATM, we performed two complementary phosphoproteomic analyses in the same samples **(Fig. 2A)**. More specifically, to gain information about direct ATM kinase targets, an ATM/ATR-motif specific antibody was employed to enrich phosphopeptides **(Supplementary Table S2)**. This revealed a total of 901 significantly dysphosphorylated peptides, with 586 peptides passing the reliability filter criteria. In ATM-kd, 104 hypophosphorylated peptides and 147 hyperphosphorylated peptides were identified (shATM vs. NT CTRL, ≥/≤ +/- 2-fold), corresponding to 74 proteins with reduced phosphorylation and 98 with increased phosphorylation **(Fig. 2B, left panel)**. These potential direct ATM targets will be represented with capital letters (P-Ser/Thr) throughout the manuscript.

**Figure 2:**
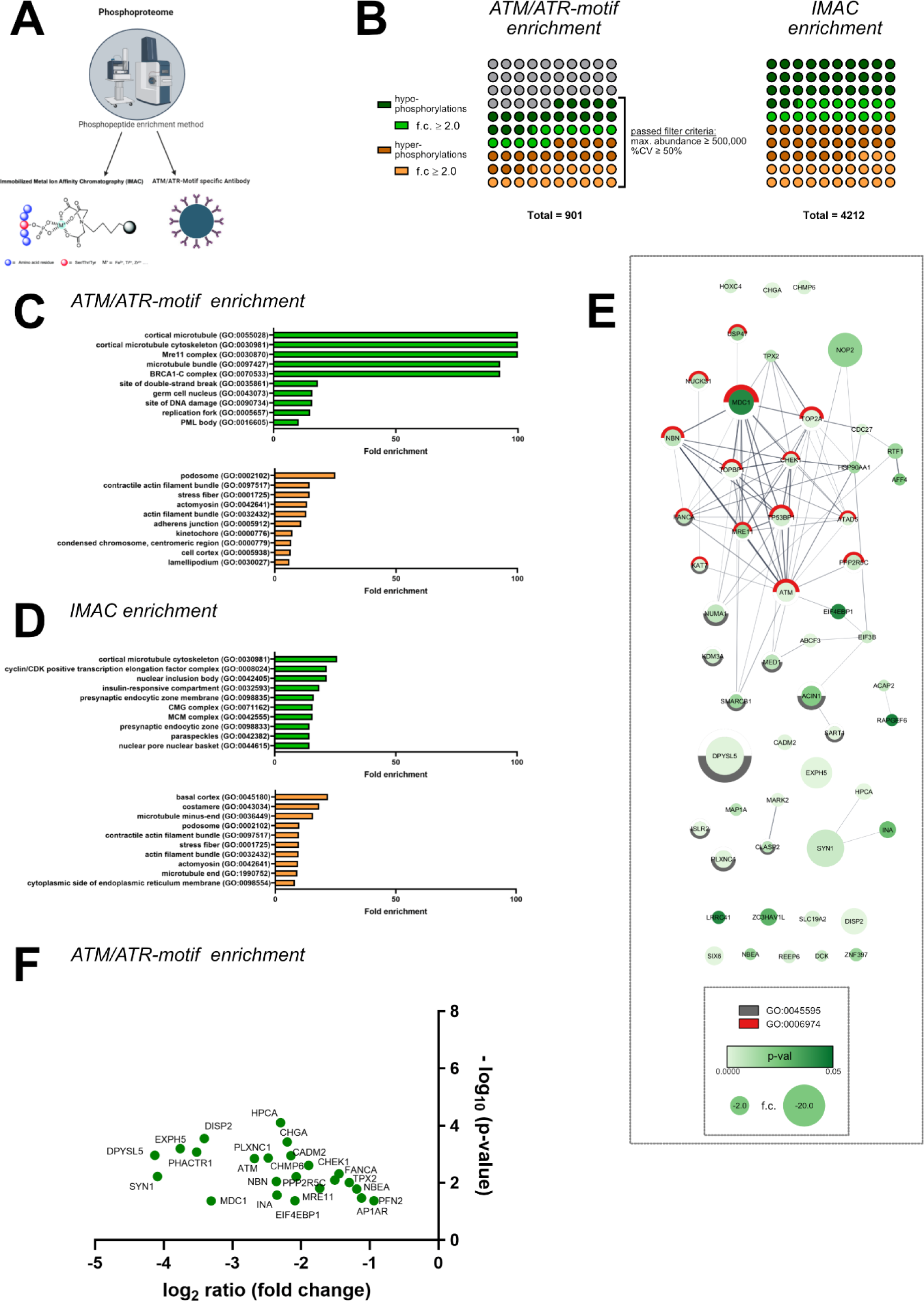
Global phosphorylome surveys in SH-SY5Y neuroblastoma cells with ATM knockdown reveal functional enrichment of microtubule cytoskeletal factors and cell differentiation processes, among the hypophosphorylations. **(A)** *In vitro* phosphoproteomics was carried out in the same samples as the previous proteome profiling. The samples were subjected to phosphopeptide enrichment via ATM/ATR-motif-specific antibody or via IMAC, each followed by mass spectrometry analysis. **(B)** Summary of results obtained from both phosphorylome surveys. **(C)** Functional enrichment of cellular components in the ATM/ATR-motif enriched screen determined by PANTHER classification system (https://pantherdb.org/) for hypophosphorylated proteins (green color) and hyperphosphorylated proteins (orange color) with fold change of at least ± 2.0 when comparing NT CTRL and shATM conditions. Biggest enrichments were found for microtubule cytoskeleton factors and DNA damage processes. **(D)** Functional enrichment of cellular components in the IMAC enriched screen determined by PANTHER classification system for hypophosphorylated proteins (green color) and hyperphosphorylated proteins (orange color) with fold change of at least ± 2.0 when comparing NT CTRL and shATM conditions. **(E)** Cytoscape network of significantly hypophosphorylated proteins from the ATM/ATR-motif specific screen with a fold change smaller that -2.0 when comparing NT CTRL and shATM conditions. Green color grading refers to significance and icon size refers to fold change (f.c.). The big cluster of proteins implicated in „cellular response to DNA damage stimulus (GO:0006974)“ is depicted with a red semicircle, proteins that are involved in „regulation of cell differentiation (GO:0045595)“ are depicted with a grey semicircle. **(F)** Volcano plot of selected hypophosphorylated proteins from the ATM/ATR-motif specific screen with fold change (calculated as log2 ratio) plotted against p-value (calculated as –log10). Abbreviations: IMAC = immobilized metal affinity chromatography.

For a general overview of phosphorylation differences, immobilized metal affinity chromatography (IMAC) enrichment of phosphopeptides was performed **(Supplementary Table S3)**, detecting a total of 4212 differentially phosphorylated peptides of which 701 were hypophosphorylations and 988 were hyperphosphorylations (shATM vs. NT CTRL, ≥/≤ +/- 2-fold), corresponding to 473 unique proteins with hypophosphorylations and 552 unique proteins with hyperphosphorylations **(Fig. 2B, right panel)**. These global phosphorylations will be represented with lowercase letters (p-Ser/Thr) throughout the manuscript.

To gain an idea about the potential pathways affected by differential phosphorylation upon ATM- kd, gene ontology enrichment analysis was performed using PANTHER for the respective unique proteins. For the ATM/ATR-specific survey **(Fig. 2C)**, we found a prominent enrichment of the cellular components “cortical microtubule (GO:0055028)”, and the “Mre11 complex (GO:0030870)” among hypophosphorylations, demonstrating that the well-known ATM interactome components acting in DDR were also detected by the obtained screen. Increased phosphorylations, which might arise from ATR-mediated phosphorylations that are equally detected by ATM/ATR-motif antibody, were especially enriched for the cellular components “podosome (GO:0002102)”, “contractile actin filament bundle (GO:0097517)” and “stress fiber (GO:0001725)”.

This was also reproduced by GO enrichment analysis of the global IMAC phosphorylome survey using PANTHER **(Fig. 2D)**, where we found the biggest enrichments for the “cortical microtubule cytoskeleton (GO:0030981)” cellular component for hypophosphorylated proteins. Among biological processes, hypophosphorylations were also enriched for “synaptic vesicle cycle (GO:0099504)” as minor effect (3.26-fold, not among the TOP10 enriched GO-terms). Considering the hyperphosphorylated proteins, enrichments were again found for “podosome (GO:0002102)” and “stress fiber (GO:0001725)” cellular components and the biological process “negative regulation of protein secretion (GO:0050709)” as a minor effect (4.75-fold, not among the TOP10 enriched GO-terms).

These phosphopeptide data indicate that the hypophosphorylations expected upon ATM deficiency are enriched in microtubule-associated cytoskeleton factors and pathways, while the hyperphosphorylations that are expected upon compensatory ATR hyperactivity are enriched in actin-rich cell extensions and contact zones.

The Cytoscape visualization tool combined with STRING functional enrichment analysis was employed to highlight interesting target proteins among the ATM/ATR-specific hypophosphorylated proteins **(Fig. 2E)**. They included factors that cluster around the biological process “cellular response to DNA damage stimulus (GO:0006974, depicted in red)”, which corresponds to the well-known canonical pathway of nuclear ATM that targets NBN, MRE11, CHEK1, MDC1 and FANCA **(Fig. 2F)**. In addition, the known cytosolic ATM phosphorylation targets EIF4EBP1, AP3B2, PPP2R5C, SYN1 and SLC2A1 were confirmed in this survey **(Fig. 2F)**. Importantly, among the novel cytoplasmic ATM kinase targets the biggest hypophosphorylation (-17.5-fold change; -4.13-fold log2 ratio shATM vs. NT CTRL; **Fig. 2E,F**) in shATM cells was found for P-Ser538 in CRMP5 (gene symbol *DPYSL5*), a protein with significantly reduced abundance **(Fig. 1C)**. In terms of functional association, this protein is implicated in “regulation of cell differentiation (GO:0045595, depicted in grey), forming a second cluster among these hypophosphorylated proteins. Previous research identified the Thr516 phosphorylation site in the tubulin binding region of CRMP5 to impact dendrite growth (Brot et al., 2014; Brot et al., 2010), highlighting that the CRMP5 protein reduction and concomitant phosphorylation deficit could impact tubulin dynamics and thereby regulate neurite morphological features. This is further supported by the high phosphorylation deficit of P-Ser978 in PLXNC1 **(Fig. 2F)**, a receptor for SEMA7A in synapses between cerebellar climbing fibres and Purkinje neurons (Uesaka et al., 2015). Strong novel phosphorylation deficits in putative direct ATM kinase targets were also evident for P-Ser1503 in EXPH5, P-Ser113 in CHGA, P-Thr130 in CHMP6, and P-Ser1007 in NBEA **(Fig. 2F)**, all of which are important regulators or components of secretory granules. Similarly strong novel phosphorylation deficits in putative direct ATM kinase targets were also documented for P-Ser1252 in DISP2, P-Ser505 in PHACTR1, P-Ser78 in INA and P-Ser252 in TPX2 **(Fig. 2F)**, all of which act as neurite extension factors.

These data demonstrate that especially proteins implicated in neuronal differentiation, regulation of axonal and microtubule integrity, and secretory proteins are potential targets for cytoplasmic ATM signaling, with the biggest impact detected for CRMP5 protein. Reduced detection of ATM autophosphorylation site P-Ser1981 in the ATM/ATR-specific screen (-6.43-fold change; -2.68-fold log2 ratio shATM vs. NT CTRL; **Fig. 2E,F**) reflects ATM depletion as positive control and supports the high quality of this dataset.

### 5.3. *In vivo* hypophosphorylations are enriched in microtubule-based transport proteins

We next compared these *in vitro* OMICS screens to *in vivo* data from cerebella of aged ATM-null (KO) mice compared to wildtype (WT) littermates. In these tissues, proteomic profiling was performed again **(Supplementary Table S4)**, together with phosphoproteome screens via ATM/ATR-motif antibody **(Supplementary Table S5)** and via IMAC **(Supplementary Table S6)**. The mutant animals developed thymomas earlier than expected and had to be sacrificed already at the age of 8 months **(Fig. 3A)**, when initial cerebellar volume reduction becomes apparent (Pietzner et al., 2013). Therefore, the anomalies were sparse, but they were consistent with *in vitro* data, revealing the molecular changes at the very beginning of pathogenesis.

**Figure 3:**
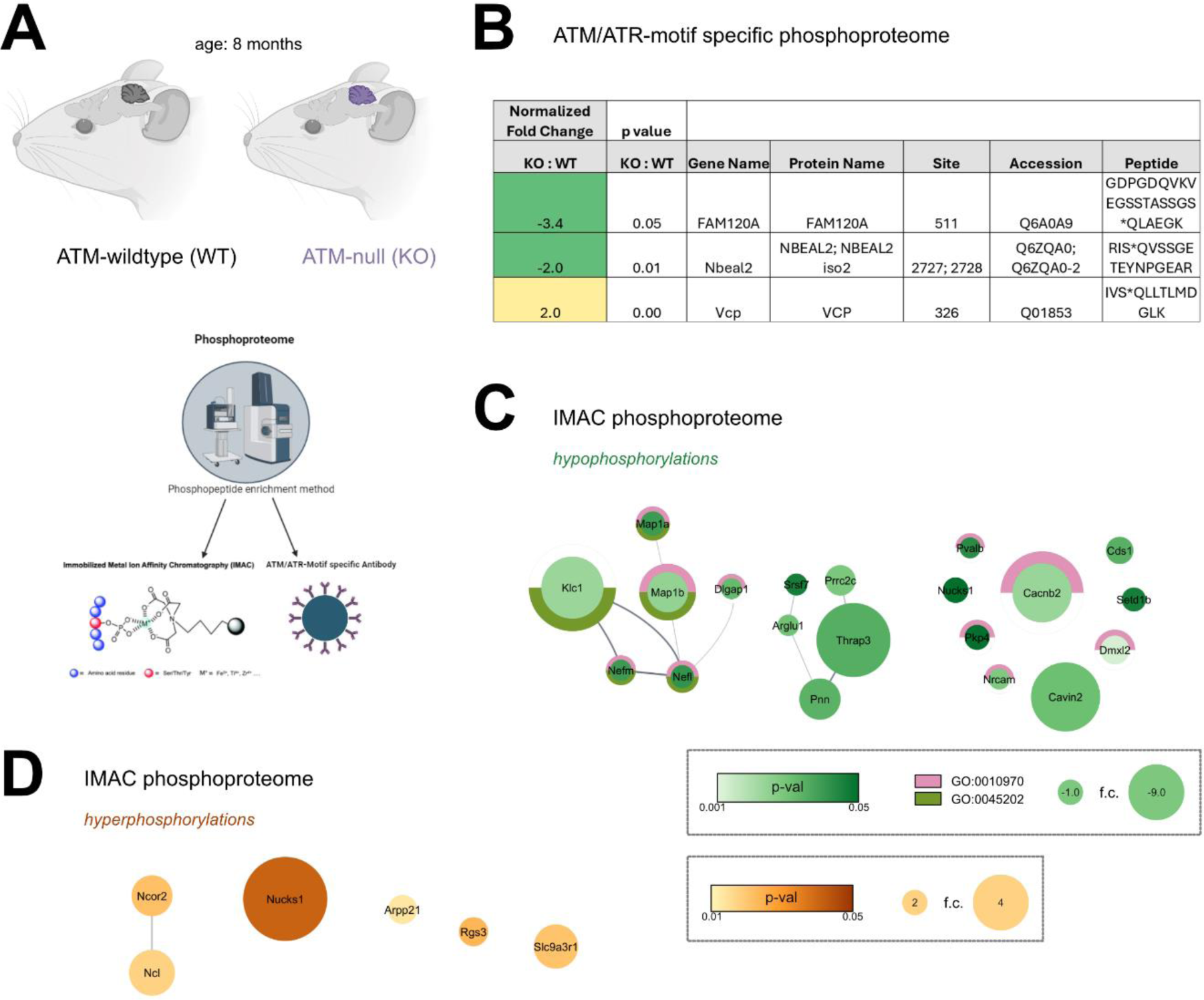
Global phosphorylome survey in cerebella of ATM-null mice demonstrates functional enrichment in microtubule-dependent transport processes, among hypophosphorylations. **(A)** Workflow of the global phosphorylome surveys *in vivo*. Cerebella of 8-month-old KO mice (ATM-null) and WT controls (wild type, 4 vs. 4) were subjected to phosphopeptide enrichment via ATM/ATR-motif-specific antibody or via IMAC each followed by mass spectrometry analysis. **(B)** Significantly dysphosphorylated peptides in cerebella determined by ATM/ATR- phosphopeptide enrichment method. **(C)** Cytoscape network of significantly hypophosphorylated proteins determined by IMAC phosphopeptide enrichment method. Color grading refers to significance and icon size refers to fold change. Proteins implicated in the cellular component „Synapse (GO:0045202)“ are depicted in mauve, proteins that are involved in the biological process „transport along microtubules (GO:0010970)“ are depicted in green. **(D)** Cytoscape network of significantly hyperphosphorylated proteins determined by IMAC phosphopeptide enrichment method. Color grading refers to significance and icon size refers to fold change.

In the proteome profile, no significant anomaly was detected. In the ATM/ATR-motif survey, three differentially phosphorylated peptides were identified **(Fig. 3B)**. Of these, two were hypophosphorylations as expected for direct ATM phosphorylation targets in KO mice. The strongest hypophosphorylation was found for P-Ser511 in the oxidative stress-responsive kinase activator FAM120A (also known as C9orf10), downstream from SRC and PI3K, with involvement in mTORC1 signaling and axonal transport (Bartolomé et al., 2015; Cho et al., 2023; Kobayashi et al., 2008; Tanaka et al., 2009). Significant hypophosphorylation was also observed for P-Ser2727 in NBEAL2 as a factor that associates with tubulin, binding to the vesicle trafficking protein LRBA, and responsible for α-granule release in platelets (Aarts et al., 2021; Cullinane et al., 2013; Delage et al., 2023; Mayer et al., 2018; Shinde et al., 2021). Converse hyperphosphorylation was documented for the known ATM-dependent stress response factor VCP (Livingstone et al., 2005) at P-Ser326. This membrane protein extraction factor plays crucial roles for endocytosis, autophagy and modulation of IGF-1 signaling (Avci and Lemberg, 2015; Bug and Meyer, 2012; Glinka et al., 2014; Ju et al., 2009; Osorio et al., 2016). Thus, trophic kinase signals that govern vesicle dynamics are altered at the earliest stage of cerebellar atrophy in the ATM-null mouse.

Integrating all cerebellar IMAC data via a STRING functional enrichment analysis with visualization by Cytoscape, the hypophosphorylated proteins exhibited overrepresentation of the cellular component „Synapse (GO:0045202)“ (depicted in mauve, **Fig. 3C**) and the biological process „transport along microtubules (GO:0010970)“ (depicted in green, **Fig. 3C**). For hyperphosphorylations, no enrichments were found **(Fig. 3D)**.

Taken together, these *in vivo* data support and refine the *in vitro* findings. They highlight the concept that ATM-deficiency triggers deficits in microtubule transport of secretory vesicles, resulting in synaptic pathology.

### 5.4. ATM knockdown induces semaphorin-CRMP5 signaling deficits *in vitro*

As functional validation of these six proteomic datasets, we examined semaphorin-CRMP5 signaling to microtubules *in vitro*, using the microtubule-stabilizing drug Taxol in the ATM-kd neuroblastoma cells **(Fig. 4A)**. The significant *ATM*-deficiency (to 45-50%) **(Fig. 4B)** with parallel *DPYSL5* transcript decrease (to ∼30%) were verified by RT-qPCR **(Fig. 4C)**, without significant modification by Taxol. Upstream, the signaling receptor *SEMA3A* mRNA was significantly induced upon ATM-kd **(Fig. 4D)**, potentially for compensatory reasons. Upregulation was observed downstream also for the CRMP5 target *TUBB3* (βIII-Tubulin) transcript, but not for *MAP2* transcript, again without Taxol modification **(Fig. 4E)**. Thus, the low abundance of CRMP5 upon deficient ATM-dependent Ser538-phosphorylation cannot be explained only via protein destabilization, but the resynthesis by transcription/translation also appeared reduced. Importantly, signaling components upstream and downstream from CRMP5 also showed altered expression.

**Figure 4:**
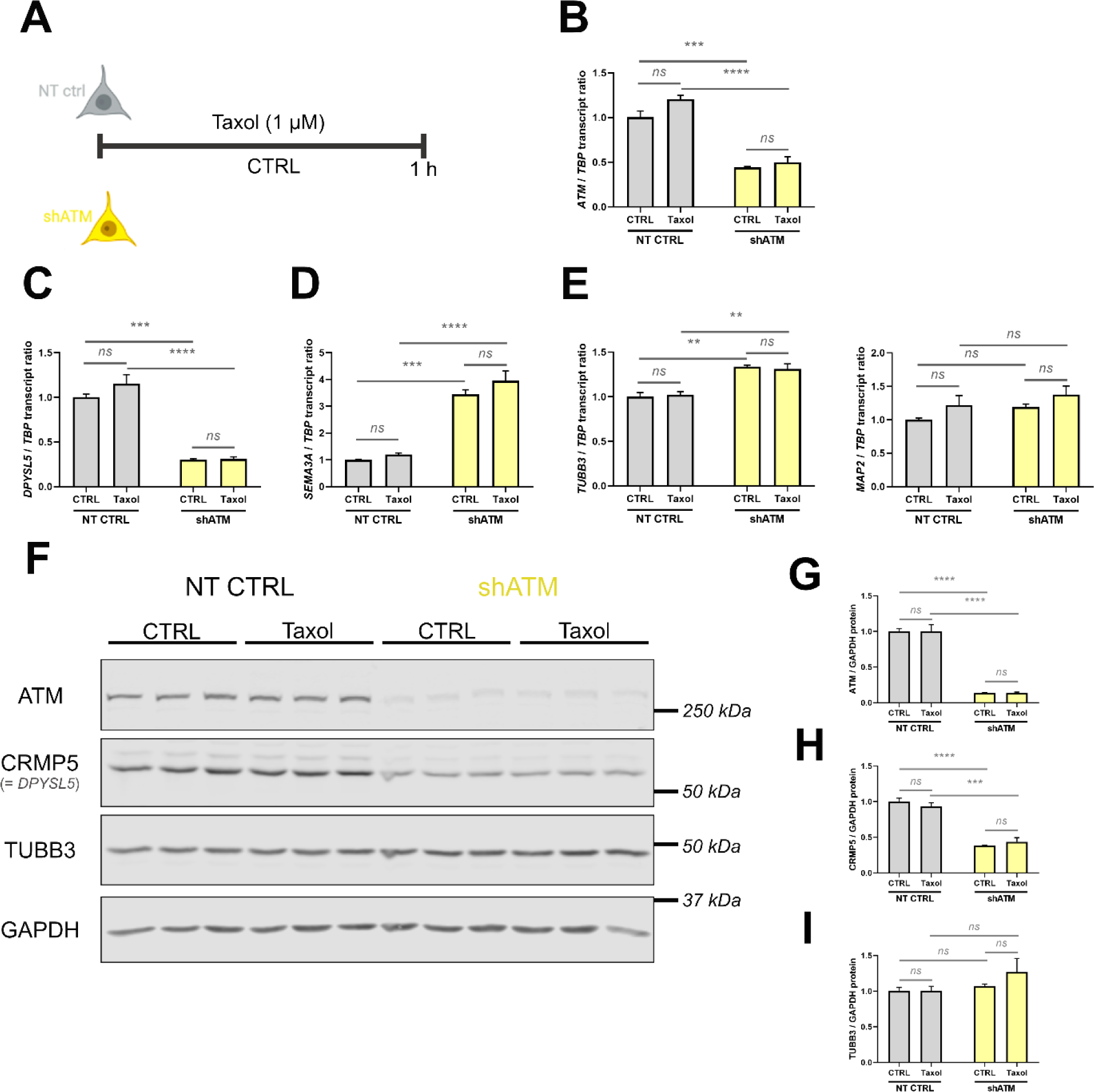
Dysregulation of the semaphorin-CRMP5 signaling axis *in vitro* upon ATM loss and microtubule-stabilizing stress. **(A)** Scheme for the *in vitro* stressor treatment using the microtubule-stabilizing drug Taxol. **(B)** Significant reduction of *ATM* transcript in shATM cells. **(C)** Significant reduction of *DPYSL5* transcript (encoding CRMP5 protein) levels in shATM cells. **(D)** Significant induction of transcript levels of the upstream signaling receptor *SEMA3A*. **(E)** Transcript levels of CRMP5 downstream signaling components *TUBB3* and *MAP2*. Transcript levels are n=3. **(F)** Immunoblots for quantification of ATM, CRMP5 and TUBB3 protein levels with GAPDH as housekeeping protein. **(G)** Densitometric quantification of ATM protein to GAPDH levels, demonstrating significant reduction of ATM protein in shATM cells. **(H)** Densitometric quantification of CRMP5 protein to GAPDH levels, with significant reduction in shATM cells. **(I)** Densitometric quantification of TUBB3 protein to GAPDH levels. Protein levels are n=3. Bar plots represent mean + SEM. Asterisks reflect significance: * = p ≤ 0.05, ** = p ≤ 0.01, *** = p ≤ 0.001, *** = p ≤ 0.0001; T = p ≤ 0.1, ns = non-significant.

Quantitative immunoblot analyses of ATM, CRMP5 and TUBB3 **(Fig. 4F)** confirmed ATM reduction to 13% **(Fig. 4G)**, CRMP5 reduction to 38-43% **(Fig. 4H)**, but unchanged TUBB3 protein amounts **(Fig. 4I)**, again without modification by Taxol administration. Immunofluorescent staining with CRMP5 and TUBB3 antibodies demonstrated CRMP5 signals to be prominent in control cells, but markedly reduced in ATM-kd cells **(Fig. S1A)**. Quantification of fluorescence intensity documented a 2-fold decrease **(Fig. S1B)**, while TUBB3 remained stable **(Fig. S1C)**.

The impact of the osmotic stressor CQ (Reichlmeir et al., 2023) caused significant *DPYSL5* and *TUBB3* transcript reduction, as opposed to *SEMA3A* transcript increase in shATM cells, while having no effect on *MAP2* transcript **(Fig. S2A)**. CRMP5 protein was diminished without apparent CQ modification, and TUBB3 protein again remained stable **(Fig. S2B-D)**.

These RT-qPCR and immunoblot results confirm ATM-dependent dysregulations in the semaphorin-CRMP5-microtubule signaling axis, with minimal effects exerted by microtubule stabilizing stress or osmotic stress.

### 5.5. ATM interacts with CRMP5 and Spastin *in vitro*

To assess the interaction between ATM and CRMP5, we performed co-immunoprecipitations in cytoplasmic fractions of SH-SY5Y neuroblastoma cells **(Fig. 5)**.

**Figure 5:**
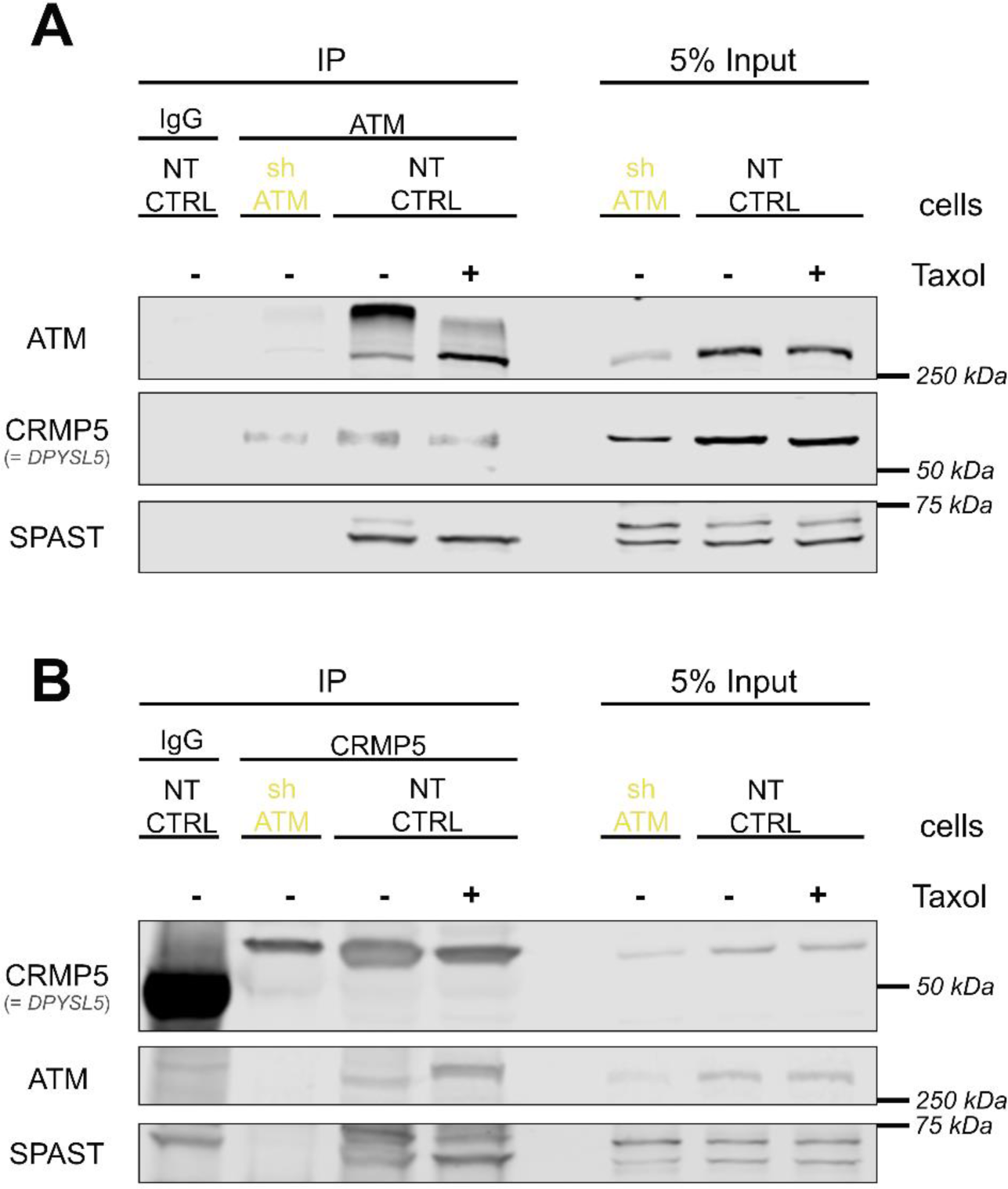
ATM interacts with CRMP5 and Spastin *in vitro*. **(A)** Immunoprecipitation of ATM as bait protein by anti-ATM (5C2) antibody in unstressed and Taxol-treated cells, with detection of CRMP5 and SPAST as prey protein. **(B)** Immunoprecipitation of CRMP5 as bait protein in unstressed and Taxol-treated cells, with detection of ATM and SPAST as prey protein. Abbreviations: SPAST = Spastin

In non-target control cells, ATM was precipitated from cytoplasmic fractions using the ATM (5C2) antibody, also upon treatment with the microtubule stressor Taxol (Fig. 5A). Importantly, CRMP5 was co-immunoprecipitated as prey protein, together with CRMP5 interactor Spastin (SPAST) especially in its small molecular weight form at 60 kDa. In smaller amount, CRMP5 co-immunoprecipitated also in shATM cells with the low residual ATM protein levels. Importantly, the co-immunoprecipitation of SPAST with ATM disappeared in shATM cells. The converse immunoprecipitation with CRMP5 antibody, followed by detection of ATM in the precipitates (Fig. 5B), detected ATM as prey protein in NT CTRL but not shATM lysates, irrespective of Taxol treatment. Upon precipitation with IgG isotype control, a generally high background with a band at a slightly higher molecular weight were confounders. A similar observation was made upon detection of SPAST as prey protein, with the co-immunoprecipitation being lost again in shATM cells. These findings suggest that cytoplasmic ATM resides in a protein complex with CRMP5 and SPAST, and that their interactions are influenced by ATM depletion.

### 5.6. Microtubules are stabilized upon ATM knockdown *in vitro*

To investigate the functional consequences of impaired CRMP5 signals for microtubule assembly (Brot et al., 2010), a commercial microtubule assay was employed. The assay performs cell lysis in a microtubule stabilization buffer, followed by gentle centrifugation to sediment large microtubule complexes attached to Golgi or nucleus as low-speed pellet (LSP), until differential ultracentrifugation of the supernatant separates soluble tubulin in the high-speed supernatant (HSS) from polymerized microtubules in the high-speed pellet (HSP), using treatment with the microtubule-stabilizer Taxol as positive control.

Immunoblotting of Tubulin, ATM and CRMP5 protein **(Fig. 6A)** in these subcellular fractions with subsequent densitometric quantification revealed that the microtubule-stabilizing drug Taxol, as well as ATM-kd, caused significant redistribution of microtubules to large complexes in the LSP fraction **(Fig. 6B)**. In contrast, treatment with CQ did not redistribute tubulin **(Fig. S3A/B)**. Free tubulin in HSS was decreased by Taxol, but this was not significant for ATM-kd. ATM was most prominently co-sedimenting with free tubulin in the HSS fraction, with a general decrease of ATM protein in shATM cells, as expected **(Fig. 6C)**, a finding that was robustly reproduced in the chloroquine stressor experiment **(Fig. S3C)**. Co-sedimentation with free tubulin in the HSS fraction was also observed for CRMP5, irrespective of Taxol **(Fig. 6D)** and CQ stressor administration **(Fig. S3D)**. These results demonstrate that microtubules are stabilized in response to ATM knockdown *in vitro*.

**Figure 6:**
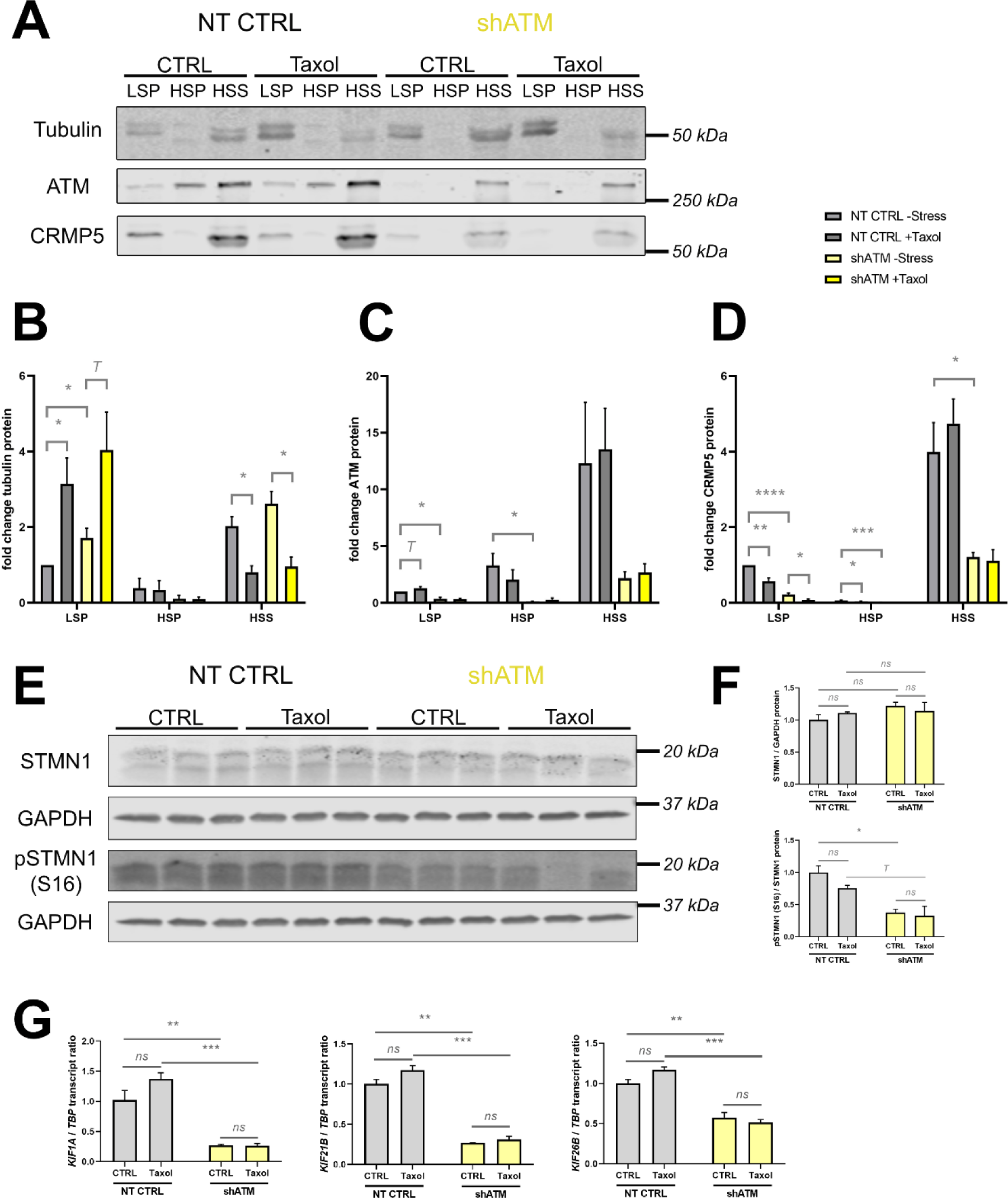
Enhanced microtubule stability *in vitro* upon ATM depletion. **(A)** Immunoblots for quantification of the microtubule content versus free tubulin and for quantification of ATM and CRMP5 protein in different fractions. **(B)** Densitometric quantification of tubulin protein in the different fractions. Tubulin protein shifted from the HSS fraction containing free tubulin towards the LSP fraction containing microtubules upon Taxol treatment but also in shATM cells. **(C)** Densitometric quantification of ATM protein in the different fractions. ATM was detected mostly in the HSS fraction and was generally reduced in shATM cells. **(D)** Densitometric quantification of CRMP5 protein in the different fractions, following a similar pattern as ATM protein. Graphs are depicted as fold changes to the unstressed NT CTRL LSP. Protein levels are n=3. **(E)** Immunoblots for quantification of STMN1 and its phosphorylation at Ser16 as a second measure for microtubule stabilization (Jourdain et al., 1997). **(F)** Densitometric quantification of STMN1 total protein levels relative to GAPDH housekeeping protein, and densitometric quantification of p-Ser16-STMN1 levels to STMN1 total protein, revealing decreased phosphorylation of STMN1 in shATM cells. Protein levels are n=3. **(G)** mRNA levels of further microtubule associated anterograde axonal transport factors, *KIF1A*, *KIF21B* and *KIF26B*. Transcript levels are n=3. Bar plots represent mean + SEM. Asterisks reflect significance: * = p ≤ 0.05, ** = p ≤ 0.01, *** = p ≤ 0.001, *** = p ≤ 0.0001; T = p ≤ 0.1, ns = non-significant. Abbreviations: LSP = low-speed pellet, HSP = high-speed pellet, HSS = high-speed supernatant.

As a second measure of microtubule stability, the STMN1 phosphorylation at Ser16 (S16) was assessed (Jourdain et al., 1997) via immunoblotting **(Fig. 6E)**. While STMN1 total protein levels where not altered upon ATM-kd **(Fig. 6F)**, significantly decreased phospho-S16 levels were observed in unstressed shATM neuroblastoma cells, while the phosphorylation reduction upon Taxol treatment did not reach significance. The ATM-dependent reduction of STMN1 S16 phosphorylation was also observed in an independent experiment using CQ **(Fig. S3E/F)**. However, a reduced phosphorylation of STMN1 at S16 was observed after CQ treatment, indicating stabilization of microtubules by osmotic stress, and contrasting with the ultracentrifugation results.

Given that axonal transport processes rely on microtubules, their affection by the microtubule stabilization in ATM-kd neuroblastoma cells was assessed. Different transport motors were analysed at mRNA level to test dependence on ATM and Taxol versus CQ stress. *KIF1A*, *KIF21B* and *KIF26B* as kinesin family members for anterograde transport were significantly reduced in shATM cells, but not altered by Taxol treatment **(Fig. 6G)**. The significant downregulation of *KIF1A*, *KIF21B* and *KIF26B* mRNA upon ATM-kd was reproduced in subsequent experiments with the stressor CQ **(Fig. S3G)**, where stressor treatment upregulated *KIF1A* and *KIF26B*, while *KIF21B* transcript showed a trend towards downregulation.

Altogether, these results indicate that ATM deficiency is associated with stabilized microtubules *in vitro,* like treatment with Taxol, while the effects of osmotic stress on microtubule dynamics are not clearcut. The data suggested that axonal microtubule-based transport could be altered.

### 5.7. ATM knockdown is associated with retracted neurites *in vitro*

Profound alterations in the cellular morphology of ATM-kd neuroblastoma cells were previously reported (Reichlmeir et al., 2023). In view of our multi-OMICS data and their functional validations, the differentiation state gains novel importance. Examining the cellular morphology with a transmitted light microscope and staining of cell nuclei using Hoechst, with subsequent measurement of neurite length, we found gross alteration of cell morphology upon ATM-kd, similar to treatment with Taxol **(Fig. 7A)**. While NT CTRL cells displayed an overall neuronal phenotype with most cells displaying neuron-like processes, ATM-kd and Taxol treatment caused round morphology and only few short processes. This was reflected by a highly significant reduction of neurite length from ∼50 µm to only ∼25 µm in Taxol-treated control cells **(Fig. 7C)**, at similar levels as seen for ATM-kd. In shATM cells, Taxol treatment could not exacerbate the morphology phenotype. In comparison, CQ treatment also shortened neurites **(Fig. 7D)**, but this effect did not reach the same magnitude as ATM knockdown and the overall morphology of SH- SY5Y neuroblastoma cells remained the same upon CQ treatment **(Fig. 7B)**.

**Figure 7:**
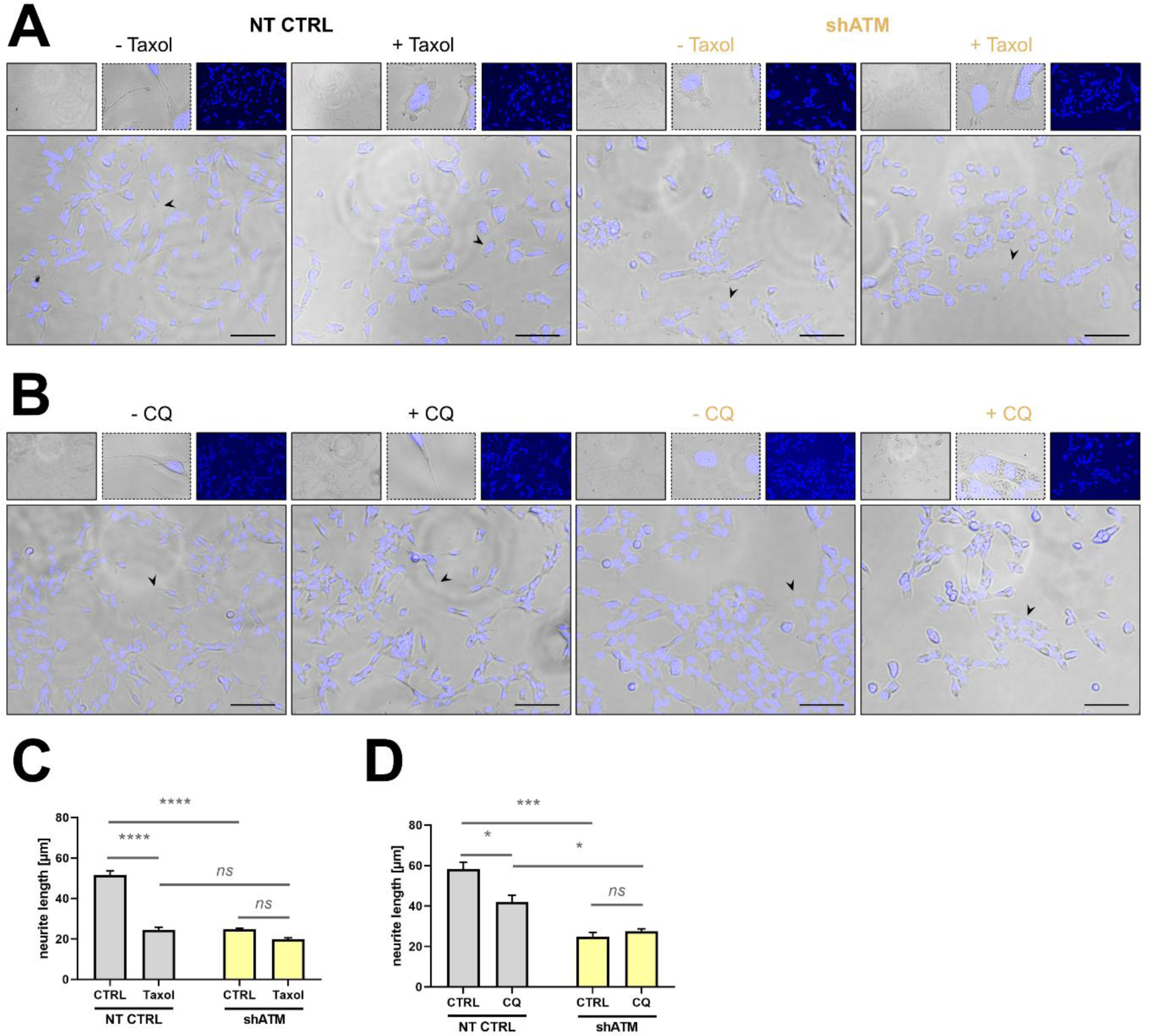
Neurite retraction induced by ATM knockdown and microtubule-stabilization *in vitro*. **(A)** Images for the analysis of SH-SY5Y morphology upon microtubule-stabilizing stress via administration of Taxol. Pictures were taken at 20x magnification with EVOS M5000 microscope in transmitted light and DAPI channel. Scale bar corresponds to 100 µm. **(B)** Images for the analysis of SH-SY5Y morphology upon osmotic stress via administration of CQ. Pictures were taken at 20x magnification with EVOS M5000 microscope in transmitted light and DAPI channel. Scale bar corresponds to 100 µm. **(C)** Quantification of neurite length in Taxol-treated cells, demonstrating significant neurite retraction, which is phenocopied in shATM cells. **(D)** Quantification of neurite length in CQ treated cells. CQ caused significant neurite retraction, but to a lower extent and without alteration of general morphology. In contrast, shATM cells displayed altered morphology and significantly shorter neurites. Neurite length was measured in three biological replicates. Bar plots represent mean + SEM. Asterisks reflect significance: * = p ≤ 0.05, ** = p ≤ 0.01, *** = p ≤ 0.001, *** = p ≤ 0.0001; T = p ≤ 0.1, ns = non-significant. Abbreviations: CQ = chloroquine.

These results suggest that altered neurite morphology might be a functional consequence of stabilized microtubules and aberrant semaphorin-CRMP5 signaling axis, which could be one reason for the selective vulnerability of neurons upon ATM deficiency.

Jointly, these results indicate that the deficits in CRMP5 signaling, probably impacting downstream microtubule dynamics via Spastin, impair neurite morphology and potentially also transport processes along microtubules for protein secretion.

## 6. Discussion

Employing global proteomics, together with phosphoproteomics of direct ATM-dependent and additional indirect IMAC changes, we successfully defined many novel molecular targets of cytoplasmic ATM in human ATM-kd neuroblastoma, and filtered the initial events of pathogenesis in 8-month-old ATM-null mouse cerebellum. It is important to emphasize how these data show excellent overlap with (i) previously published proteome profiles of A-T patient cerebella (Lee et al., 2021), (ii) A-T patient cerebrospinal fluid (CSF) (Canet-Pons et al., 2018), (iii) 12-month-old ATM-null mouse cerebellar transcriptome profiles (Reichlmeir et al., 2023), (iv) reports on ATM association with vesicles and SYN1 (Li et al., 2009), (v) with β-adaptin / AP3B2 (Lim et al., 1998), (vi) with a PP2A regulatory subunit (PPP2R5C) (Shouse et al., 2011), (vii) with ribosomal EIF4EBP1 (Mallory and Petes, 2000; Yang and Kastan, 2000), and (viii) ATM roles for NBN-MRE11-MDC1- CHEK1 in the nucleus (Lee and Dunphy, 2013; Paull, 2015). To define novel ATM-dependent proteins and pathways, in this discussion we will first consider the very strongest dysregulations in all OMICS profiles in their consistency through species and cell types, attempting to compose a plausible detailed scenario that underlies the neural atrophy process in A-T. As a broad initial concept, it seems that ATM loss leads to unrepaired DNA damage in the nucleus, and in parallel to a cytoplasmic restriction of translation, vesicular secretion, and neurite outgrowth. Mechanistically, these pathway effects appear mainly mediated by ATM controlling EIF4EBP1 at P-Ser112/P-Ser96, CHGA at P-Ser113 (and EXPH5 at P-Ser1503), as well as CRMP5 at P-Ser538. A compilation of all available evidence for each claim is provided in the first two chapters of the discussion, where the impact of these factors on their subcellular context is explored in molecular detail. The subsequent chapters of discussion are focused on the functional consequences on neurite growth, and the involvement of novel cytoplasmic ATM kinase targets in the pathogenesis of A-T and other neurodegenerative processes.

### 6.1. Cytoplasmic ATM impact on vesicular secretion

#### OMICS profiling suggests altered protein composition of small synaptic vesicles (SSV)

Cytosolic ATM is known to be associated with vesicles, also in synapses where it co-purifies with SYN1 (Li et al., 2009), and its dysfunction impairs the ability of presynapses to sustain intense electrophysiological signaling (Vail et al., 2016). In our proteome profiles, reduced SYN1 phosphorylation at P-Thr448/Ser449 was among the strongest effects, suggesting abnormal SYN1 dissociation from completely filled vesicles when they move from the reserve pool to the readily releasable pool (Sansevrino et al., 2023). If the neuroblastoma findings correctly model the events in A-T synapses of the cerebellum, then ATM seems to control the precursors of small synaptic vesicles (SSV) during their loading. The prominent converse transcript dysregulation of VGLUT1 versus VGLUT2 (*Slc17a7* and *Slc17a6*) in ATM-null mouse cerebella also suggested abnormal neurotransmitter loading of SSV precursors (Reichlmeir et al., 2023). Selective deficits in the ATM- kd neuroblastoma cell proteome were apparent in the abundance of RAB3C, RPH3A and STX1, suggesting reduced vesicle release and mobilization capacity (Fischer von Mollard et al., 1994; Foletti et al., 2001; Salazar Lázaro et al., 2024). The proteome profile also detected a reduced abundance of β-SNAP (NAPB) and SNAP25 suggesting abnormal vesicle priming and fusion (Burgalossi et al., 2010). In contrast, SYPL1, SYT4 and SYT9 were accumulated, demonstrating a perhaps compensatory effort in neuropeptide pathways (Brooks et al., 2000; Haass et al., 1996; Seibert et al., 2023; Zhang et al., 2011). Neurotransmission excitability appeared altered in view of reduced P-HPCA at Ser191 (Amici et al., 2009; Helassa et al., 2017), and p-ITPR1 in shATM cells. In good agreement, decreased *Itpr1* expression was observed in ATM-null cerebella (Reichlmeir et al., 2023). The decrease of ITPR1 expression is strong enough to predispose patients for a clinical manifestation of cerebellar ataxia (Reichlmeir et al., 2023; Tada et al., 2016).

#### Secretory granules are probably deficient

The influence of ATM on neuronal large dense core vesicles (LDCV) (outside synapses known as secretory granules) appeared much stronger and constitutive, confirming the previous evidence from ATM-null mouse cerebellar transcriptome profiling that neuropeptide pathways exhibit a systematic distortion (Reichlmeir et al., 2023). All these granules form upon production and aggregation of their common cargo, the chromogranin and secretogranin protein family, which enable dense loading despite osmotic pressure, to then be filled with the specific signaling molecules for each granule type (Kim et al., 2006). A main finding of our *in vitro* proteome profiles is the confirmation that neuropeptide signals and trophic secretion show impairment in general manner, with a massively reduced abundance of chromogranins CHGA and CHGB to 18% and 47%, respectively. There was a concomitant loss of the ATM-dependent P-Ser113-CHGA peptide to 22% opposed by an increase of the P-Ser259-CHGB peptide to 210%. Within this same protein family, IMAC phosphorylation changes were observed with increased p-Ser391-CHGB, elevated secretogranin p-Ser205-SCG5 peptide and neurosecretory trophic p-Ser360-VGF. Also the protein amounts of constitutive chromaffin granule components such as CYB561 and DBH (Winkler and Fischer-Colbrie, 1998) were reduced to 26% and to 9%, respectively. Importantly, CHGB levels are also low in patient CSF, where a markedly reduced protein secretion may contribute to low osmotic pressure in the CSF of A-T patients and thus explain the age-associated increase of the osmotic regulator albumin inside the blood-brain barrier (Woelke et al., 2021). An organism-wide decrease of vesicular protein secretion might impact the osmotic pressure also in blood, contributing to altered vascular tone, capillary dilatation, and elevated blood serum levels of the osmotic regulator AFP in A-T (Stray-Pedersen et al., 2007).

#### CHGA levels in patient blood may be a useful biomarker

These first principal findings of CHGA deficiency and decreased CHGA secretion may have clinical phenotype and diagnosis consequences, since chromogranins in the extracellular space and blood stream get cleaved proteolytically into various peptides, which have relevant roles for vascular and metabolic regulation (Garg et al., 2023; Sahu et al., 2019; Troger et al., 2017). As a CHGA cleavage product, Serpinin modulates transcription of PN-1 (*Serpine2*) that controls LDCV biogenesis (Koshimizu et al., 2010). Indeed, the transcript levels of the gene family member *Serpine3* is abnormal in 12-month-old ATM-null mouse cerebella. Increased blood CHGA is a specific and sensitive biomarker of neuroendocrine tumors (Baudin et al., 2001; Dam et al., 2020), so it may be beneficial to assess if the putative reduction of CHGA blood levels in A-T patients shows usefulness to monitor the neurodegenerative process.

#### Neuropeptide signaling shows widespread anomaly

This study observed systematic neuropeptide anomalies as expected for an LDCV release deficit (Sirkis et al., 2013). As one example, the deficiency of CHGA is known to suppress IGF signals (Zhang et al., 2019), and indeed deficient IGF-1 levels are a feature of A-T patient blood (Nissenkorn et al., 2016). At the same time, IGF-1 enhances the secretion of CHGA (Mergler et al., 2005; Münzberg et al., 2015; Wichert et al., 2000) and of other neuropeptides (Lee et al., 1999), so it would be difficult to determine if CHGA or IGF-1 is the upstream molecule. The observation that the absence of ATM massively downregulates the ATM/ATR phosphosite CHGA-Ser113, but none in the IGF-1 pathway, suggests that CHGA phosphorylation deficits and CHGA depletion are the primary events. Given that ATM associates with the outside of vesicles, whereas CHGA localizes inside vesicles, suggests that any CHGA phosphorylation by ATM would have to occur shortly after translation at the rough endoplasmic reticulum, before CHGA is translocated to the ER lumen, and subsequently triggers secretory vesicle budding at the trans-Golgi apparatus. Also the abnormal insulin-glucagon signaling and further endocrinology phenotypes in A-T may be explained via deficient secretion of an abnormally low number of CHGA or CHGB containing vesicles that were filled with various neuropeptides (Andrade et al., 2015; Bar et al., 1978; Nissenkorn et al., 2016; Spears et al., 2017; Weiss et al., 2016). A-T patient blood contains abnormal FSH levels. FSH biosynthesis is enhanced by activin-dimers and suppressed by inhibin- dimers, which are secreted from the pituitary gland. Inhibin secretion is diminished by hypothalamic GnRH but stimulated by IGF-1, while activin production is repressed by follistatin (Farnworth, 1995; Jin and Yang, 2014). Therefore, it may also be relevant that two follistatin- homologous proteins, FSTL1 and SPARC (Chen et al., 2020), were accumulated in ATM-kd neuroblastoma cells, and that patient CSF showed high FSTL5 but low SPARC. Hypothalamic parvicellular GnRH neurons express galanin (GAL) to modulate GnRH, and the anterior pituitary cells also express GAL to modulate GH, TSH and ACTH, so the low GAL abundance in ATM-kd neuroblastoma cells may also be meaningful. Furthermore, the expression of GAL in the magnocellular hypothalamic neurons acts as modulator of osmoregulation throughout the organism (Mechenthaler, 2008). Selectively in the cerebellum, GAL has a crucial role as an activator of cerebellar granule neuron migration (Komuro et al., 2021), a developmental phenotype that is affected in Nijmegen Breakage Syndrome, a disorder similar to A-T (Zhou et al., 2020). In the ATM-kd IMAC profile, massive reductions of neuropeptide receptor phosphorylations stood out for p-OPRM1 and p-NPY2R, which is in line with the 12-month-old ATM-null mouse cerebellar transcriptome where the strongest splicing change was observed for *Oprm1* (Reichlmeir et al., 2023). Neuropeptide signaling is also central in the cellular responses to irradiation or UVB exposure, and indeed various neuropeptide signaling components of the somatostatin, tachykinin, neurotensin, endothelin, vasohibin, encephalin, opioid mu and kappa3 pathways were found to be transcriptionally upregulated in the ATM-null mouse cerebellum (Reichlmeir et al., 2023). Overall, a big variety of neuropeptide pathways exhibited abnormal signaling upon ATM deficiency, and they may all be consequences of secretory vesicle deficits in chromaffin cells in the adrenal medulla or in neuroendocrine hypothalamic cells.

#### Altered neurotrophin signaling

Downstream from the secretion granule deficit, not only neuropeptides were abnormal but also various trophic factors showed prominent dysregulation. In the ATM-kd proteome, the known IGF- 1 deficit of A-T patient blood was reflected by high levels of PAPPA while IGFBPL1 was low (ATM-null mouse cerebella exhibit high *Igf1* transcript) (Gaidamauskas et al., 2013). In the ATM-kd cells, the trophic factor PTN accumulated, while its target receptor ALK and the receptor tyrosine kinases RET and LYN showed low abundance, and their intracellular downstream effector GAP43 was elevated (Yanagisawa et al., 2010). Similarly, HGF accumulated and p-HGF was elevated in the IMAC profile. In the ATM-kd proteome, the trophic receptors GFRA3 and PDGFRA showed outstandingly low levels (ATM-null cerebella exhibit high *Gfra1* and *Gfra2*, but low *Pdgfrl1* transcript), whereas SORCS1 and TNFRSF12A as receptor of the synaptic inhibitor TWEAK (Nagy et al., 2021) were strongly increased (ATM-null cerebella exhibiting elevated *Sorcs1*, *Tnfrsf13C* and *Tnfrsf21* transcripts). These observations suggest impaired trophic regulation. Thus, it may be useful to investigate, whether the secretion of specific signaling ligands from immunological synapses and the abundance of their receptors is also widely altered in A-T, as suggested by preliminary isolated reports (Covino et al., 2024; Zielen et al., 2021).

#### Vesicle traffic appears abnormal between trans-Golgi and plasma membrane

To address the question at which stage of vesicle maturation and trafficking the first anomalies are observed, we compiled all dysregulations within Rab pathways in the three proteome profiles. Reduced protein abundance was found for RAB3C, RAB33A, RAB39A and RAB28, while increased amounts existed for RAB27A, RAB8B, RAB3B, RAB31 and RAB11FIP1. ATM/ATR specific phosphorylation was increased for RABEP2, RAB3GAP2 and RAB32, while IMAC phosphopeptides were reduced for RAB13, RAB10 and RAB35, but increased for RAB3IL1, RAB3GAP1, RAB11FIP1, RAB11FIP5 and RABEP2. These observations suggest that endosome dynamics is principally affected, while ER-to-Golgi trafficking problems are not evident. Primary candidates for ATM-dependent endosome control, with predicted ATM-phosphorylation sites that are massively reduced in ATM-kd neuroblastoma cells, comprise the RAB27B effector EXPH5 at P-Ser1503 (Bare et al., 2021), the multivesicular body and ESCRT-III component CHMP6 at P-Thr130 (Yorikawa et al., 2005), and particularly RAB4-interacting neurobeachins (Gromova et al., 2018) as regulators of secretory granules that contain trophic factors. Protein abundance was reduced for NBEAL1, a disease protein for the motor neuron degeneration ALS2 (Hadano et al., 2001). In ATM-kd neuroblastoma cells, ATM/ATR mediated P-Ser1007-NBEA was reduced. In 8-month-old ATM-null mouse cerebellum, P-Ser2727-NBEAL2 decrease was an initial pathogenesis event. NBEAL2 is important for dense granule formation in platelets (Cullinane et al., 2013), and platelet granules and neuronal granules are known to share similar mechanisms (Goubau et al., 2013). Furthermore, the observed RAB33A deficit would direct LDCV trafficking away from secretion towards autophago-lysosome degradation, and indeed elevated autophagy was documented by the high abundance of DEPTOR, SQSTM1, HSPB8 and BAG3 (Nivon et al., 2016), contrasting with lowered UBQLN2 and SLC2A1 abundance presumably via autophagic degradation (Cheng et al., 2020). In view of these observations, it is conceivable that abnormal phagocytosis of microbes or their components, such as zymosan, makes a relevant contribution to the immune deficit of A-T patients, as reported for 4 A-T cases (Forte et al., 2005). Further evidence for deficient trans-Golgi to endosome trafficking can be found in the proteome profile of ATM-kd neuroblastoma cells, where the endosomal anterior pole migration factor ASTN2 and the endocytosis AP3B2 (β-NAP) adaptor protein subunit abundance was substantially diminished, accompanied by a parallel reduction of ATM/ATR specific phosphorylation of AP1AR regulatory protein. The adaptor protein subunits β-adaptin and β-NAP are known interactors of ATM (Lim et al., 1998). AP3 coating dysfunction would interfere with synaptic vesicle budding from endosomes (Faúndez et al., 1998) and is a cause for autoimmune mediated cerebellar ataxia (Jarius and Wildemann, 2015; Newman et al., 1995) as well as early onset epileptic encephalopathy (Assoum et al., 2016).

#### The vesicle secretion deficit affects specific neural adhesion proteins and extracellular matrix enzymes

A comparison of the proteome profile of ATM-kd neuroblastoma cells with the CSF proteome of A-T patients always detected accumulations within cells that contrast with decreased extracellular levels. This general pattern included the adhesion factors CDH2, EPDR1 and MCAM, as well as the lysosomal-extracellular glycoprotein-processing enzymes NEU1, MANBA and CTBS. Importantly, the secretion impairment also affected a pathway that is very well known to control cerebellar ataxia: The elevated cellular abundance of RELN and its alternative receptor ITGA3 (Rodriguez et al., 2000) in ATM-kd cells contrasts with its low levels in patient CSF. The only exception from this rule was the low abundance of the matrix repair enzyme MMP2 (Verslegers et al., 2013), both in the ATM-kd neuroblastoma cells and in the patient CSF. Overall, neural signaling, trophism, and attachment were altered as consequence of the presumptive secretion deficit in ATM-kd neuroblastoma cells, in very good agreement with ATM-null mouse cerebellum and A-T patient CSF data.

### 6.2. Cytoplasmic ATM impact on synaptic pruning and cytoskeletal regulation Growth cone chemotaxis

In view of altered neuropeptides, neurotrophins, and matrix cues, it seems logical that synaptic connections become unstable and that pathfinding for new cell contacts is impaired. As the second principal finding of this study, and as completely novel insight, ATM dysfunction controls this collapse and motility of growth cones via strong CRMP5 depletion in ATM-kd neuroblastoma cells, a finding analogous to the previously observed deficiency of CRMP3 (gene symbol *Dpysl4*) transcript in the cerebella of ATM-null mice and A-T patients (Reichlmeir et al., 2023). The P- Ser538 phosphopeptide in CRMP5 is predicted to be under ATM control, is detected by the ATM/ATR-motif antibody, and may control the stability and interactions of CRMP5. This phosphosite is located near the C-terminus in a region known to bind the neurodegenerative disease protein SPAST (Hazan et al., 1999; Ji et al., 2018). Also CRMP1-4 were shown to regulate growth cone dynamics via the microtubule and actin cytoskeleton (Aylsworth et al., 2009; Fukata et al., 2002; Ji et al., 2021; Khazaei et al., 2014; Lin et al., 2011). The crucial role of CRMP5 in neurodegenerative processes is substantiated by genetic and autoimmune clinical evidence. Missense variants in the *DPYSL5* gene are associated with neurodevelopmental disorders such as Ritscher-Schinzel syndrome, with prominent cerebellar hypoplasia and corpus callosum agenesis, resulting in ataxia and strabismus (Jeanne et al., 2021). Mechanistically, the p.E41K and p.G47R variants impair the interaction of CRMP5 protein with the microtubule binding protein MAP2 and the neuronal βIII-tubulin (TUBB3) (Jeanne et al., 2021). Since interaction of CRMP5 with MAP2 and TUBB3 is also regulated by phosphorylation in the tubulin binding region (Brot et al., 2014), the control of this complex assembly may be the responsibility of CRMP5. In our co-immunoprecipitation experiments we indeed confirmed interaction of ATM with CRMP5 and SPAST. CRMP4 mutations are associated with motor neuron degeneration in Amyotrophic Lateral Sclerosis (Blasco et al., 2013; Maimon et al., 2021). In cancer patients, the growth of lung tumors or thymomas can be complicated by the appearance of paraneoplastic autoantibodies against CRMP5, CRMP4 or CRMP3, which trigger of neurodegenerative processes, affecting preferentially the cerebellum in a quarter of cases (Guasp et al., 2021; Knudsen et al., 2007; Yu et al., 2001).

CRMPs were first discovered (Goshima et al., 1995) as homologs of the *Caenorhabditis elegans* unc-33 axonal guidance protein. These proteins are the intracellular signaling mediators of secreted extracellular guidance cues such as collapsin / semaphorin3A (SEMA3A) (Luo et al., 1993) and ephrins (Arimura et al., 2005; Hou, 2020; Schmidt and Strittmatter, 2007). It is therefore relevant that ATM-kd neuroblastoma cells showed high SEMA3A/C (versus low SEMA7A in patient CSF) and dysregulation of their co-receptors (PLXNA2, PLXNA4 and NRP1 being increased in the proteome, P-PLXNC1 decreased in ATM/ATR phosphorylome). Within the ephrin pathway, EPHA2 was elevated (in ATM-null cerebellar transcriptome *Epha4/5* up, *Epha3* down), while EPHB2 was decreased (in ATM-null cerebellum *Efnb3* transcript up). Growth cone motility and axon pruning critically depend on dynamic microtubules, since microtubule catastrophe and rescue are both required for axonal outgrowth and path finding (Tanaka and Kirschner, 1991). Mechanistically, it was demonstrated for CRMP2 that it impacts axonal growth via a direct interaction with tubulin heterodimers to activate microtubule assembly and thereby increase neurite formation (Fukata et al., 2002). Similarly, CRMP5 directly interacts with tubulin via a C-terminal tubulin binding domain, and this interaction is important for the inhibition of tubulin polymerization into microtubules, thereby promoting inhibition of neurite outgrowth (Brot et al., 2010). CRMP5 is regulated by phosphorylation at Thr516, which is located in the tubulin-binding domain. This phosphorylation is mediated by glycogen synthase kinase 3β (GSK-3β), facilitating CRMP5 interaction with tubulin, thereby disrupting tubulin polymerization and efficient counteraction of CRMP2-mediated promotion of dendrite outgrowth (Brot et al., 2014). Via an OMICS-based approach, it was also demonstrated that ATM interacts with several cytoskeletal proteins and microtubule components in HeLa cells in the presence or absence of DNA damaging stress by ionizing radiation (IR), and in fibroblasts upon cytoskeletal stress (Bastianello et al., 2023). Overall, axon guidance and neurite growth signaling is abnormal upon ATM-loss, potentially via the altered CRMP5-axis.

### 6.3. Cytoplasmic ATM impact on neurite extension

#### Microtubule and actomyosin cytoskeleton providing vesicles and motility for neurites

The ATM effect via CRMP5 on microtubules was also supported by the strong depletion of postsynaptic scaffold and microtubule interactor DLGAP5, the reduction of P-Thr66-DLGAP5, and the increase of p-DLGAP4 in ATM-kd neuroblastoma cells. Also during the initial anomalies in ATM-null cerebella, p-DLGAP1 reduction was identified. Furthermore, the microtubule dysregulation was supported by a strong deficiency of the neural migration regulator DCX and a reduced abundance of the microtubule-associated protein MAP2. In addition, several ATM/ATR phosphopeptides of MAP1A were decreased, and IMAC phosphopeptide deficiency upon ATM-kd was found for MAP1A, MAP1B, MAP2, MAP7 and MAPT. Regarding the ATM effect on actomyosin, elevated levels of TAGLN2 (also in patient CSF) and reduced PFN2 were documented in the proteome, accompanied by a strong increase of P-VCL and a reduction of P-PHACTR1 in the ATM/ATR profile. Not only the actin cytoskeleton of filopodia/lamellipodia and the microtubules are affected, but also the cytoskeletal intermediate filament alpha-internexin (INA) upon ATM/ATR screening of neuroblastoma cells showed a massive reduction of the P-Ser78 phosphopeptide, while VIM, LMNA, SYNM protein and P-LMNA, P-SYNM and P-NES were increased in ATM/ATR surveys. Furthermore, in ATM-kd neuroblastoma cells, the intermediate neurofilament isoforms NEFL, NEFM and NEFH showed strong downregulation, with a corresponding decrease of IMAC phosphopeptides for NEFM and NEFH, while the early-stage disease effect in ATM-null mouse cerebellum included the reduction of axonal p-NFL and p-NFM, as well as axo-dendritic p-MAP1A and p-MAP1B. Indeed, the neuronal loss of NFL is an early and progressive feature of A-T patients (Donath et al., 2021). Overall, the cytoskeletal data indicate growth cones, axons and dendrites to be affected.

Also the ATM-null mouse cerebellar *in vivo* data indicate that the microtubule cytoskeleton is abnormal, with P-Ser511-FAM120A hypophosphorylation. FAM120A was identified as a component of mRNA transport granules along microtubules in neurons (Kobayashi et al., 2008; Ohashi et al., 2000). Previous studies already suggested ATM to be implicated in cytoskeletal regulatory processes (Kreis et al., 2019; Kreis et al., 2013). This is further supported by the notion that the SQ/TQ cluster domain (SCD), which is an accumulation of the ATM/ATR phosphorylation motif within a specific region, is overrepresented in proteins not only involved in the DNA damage response but also in proteins involved in the actin cytoskeleton, in vesicle trafficking pathways and the microtubule cytoskeleton (Cara et al., 2016; Kim et al., 1999; Traven and Heierhorst, 2005). Only very recently, it has been found that ATM is signaling via cytoskeletal pathways in response to mechanical stress probably for recovery from mechanical stress *in vitro* (Bastianello et al., 2023). This demonstrates that cytoplasmic ATM is clearly implicated in the maintenance of cytoskeletal structures and stress response pathways, in particular microtubule dynamics pathways for neuronal differentiation as identified in the present study. Dynamic microtubules are especially important for neurons regarding their morphogenesis, axon guidance, dendrite formation, growth cone motility and axonal transport (Lasser et al., 2018). We observed profound differences in microtubule stability upon ATM-deficiency, with microtubules being significantly stabilized, similar to the effects generated by the stressor Taxol. Various neurodegenerative diseases are associated with altered microtubule dynamics. The microtubule-associated protein tau, for example, is a regulator of microtubule stabilization, assembly and bundling. The pathological hyperphosphorylation of Tau in Alzheimer’s disease is associated with microtubule destabilization (Del Alonso et al., 2006; Noble et al., 2013; Prezel et al., 2018). Conversely in hereditary spastic paraplegia (HSP), mutations in the Spastin (*SPG4*) gene cause protein loss-of-function and stabilized microtubules as a consequence (Evans et al., 2005; Hazan et al., 1999). Spastin (SPAST) also interacts with CRMP5 to regulate microtubule dynamics, thereby promoting neurite outgrowth (Ji et al., 2018). This interaction occurs between the SPAST N-terminal region (residues 270-328) and CRMP5 C-terminal region (residues 472-564) (Ji et al., 2018), meaning that the CRMP5 binding site in SPAST is located near the p-Ser268 residue, where phosphorylation decreases upon ATM-kd. Similarly, the CRMP5 P-Ser538 site is within its SPAST binding region. SPAST phosphorylation at Ser-268, a HIPK2 target during cytokinesis, is necessary for abscission and midbody localization, preventing SPAST degradation by NEDDylation (Pisciottani et al., 2019; Sardina et al., 2020). SPAST is also involved in early secretory pathways and endosomes, predominantly as the 60 kDa isoform (Connell et al., 2009). There, SPAST may be recruited to endosomes at axonal branches to provide cargos for growth (Connell et al., 2009). Our co-immunoprecipitation data showed that the 60 kDa SPAST interacts with CRMP5 in control cells, but not in ATM-kd cells. This indicates that CRMP5 and SPAST might work together to regulate microtubules and neurite growth, and dysfunction in this interaction could explain enhanced microtubule stability and neurite retraction in ATM-kd cells. This synergy is also important in mouse oocyte meiosis, where CRMP5 deletion causes spindle issues that can be rescued by Spastin overexpression (Jin et al., 2021). Female ataxia telangiectasia patients lack oocytes due to meiotic degeneration (Nissenkorn et al., 2016; Xu et al., 1996), a phenomenon possibly linked to ATM, SPAST, and CRMP5 dysfunction. To evaluate the observed microtubule stabilization upon ATM loss by a second method, the analysis of STMN1 phosphorylation at Ser16 was employed. This experiment confirmed that ATM-kd affects the stability of microtubules. The role of STMN1 phosphorylation in modulating its microtubule-destabilizing activity has been a subject of debate, with opposing findings reported in the literature. It has been suggested that phosphorylation of STMN1 leads to a reduction in its microtubule-destabilizing activity (Horwitz et al., 1997; Marklund et al., 1996), while other studies found contrasting evidence, indicating that STMN1 phosphorylation activates and induces depolymerizing activity (Belmont and Mitchison, 1996; Jourdain et al., 1997). Furthermore, STMN1 has further regulatory phosphorylation sites at Ser25 and Ser38, which are all important for its activity towards microtubules (Di Paolo et al., 1997). Given that tubulin mRNA levels are regulated by autoinhibition in response to microtubule stabilization and destabilization, the increased *TUBB3* mRNA in cells with ATM-kd provided further support for microtubule stabilization (Gasic, 2022; Gasic et al., 2019; Lin et al., 2020). Overall, our data demonstrate that microtubules are stabilized *in vitro* upon ATM loss, as a potential downstream effect of dysfunctional CRMP5 signaling axis. Morphologically, the stabilization of microtubules by Taxol is already known to impair neurite branching and cause growth inhibition and abnormal neurite directionality, while increasing neurite thickness (Chuckowree and Vickers, 2003; Ferrari-Toninelli et al., 2008; Letourneau et al., 1986). This is in line with the drastic morphological differences observed *in vitro* upon ATM loss (Reichlmeir et al., 2023), which can be promoted by the application of Taxol alone. Neurites are virtually absent upon ATM-kd, where SH- SY5Y neuroblastoma cells acquire a round shape with several very small protrusions.

Since dysregulations in pathways of osmoregulation were observed by transcriptomic *analysis* in cerebella of 12-months old ATM-null mice, it was relevant to study how osmotic stress impacts microtubule cytoskeletal dynamics *in vitro*. As stressor in these experiments, we applied chloroquine, originally described as anti-malaria drug and inhibitor of autophagy (Mauthe et al., 2018). Since chloroquine is a cationic drug, it accumulates in acidic compartments and draws water by osmotic mechanisms, leading to vacuolar response in the cell (Marceau et al., 2012). Chloroquine itself, as well as hypotonic stress, is known to activate ATM, demonstrated by elevated phosphorylation of S1981 autophosphorylation site (Bakkenist and Kastan, 2003; Qian et al., 2018; Reichlmeir et al., 2023). Apart from that, chloroquine administration disrupts the actin cytoskeleton in podocytes, thereby influencing cell motility (Kang et al., 2020). This demonstrates that microtubule signaling could be an important pathway for ATM in the cytoplasm potentially also in response to osmotic stress, as we demonstrated CRMP5-microtubule signaling axis to be also impacted after CQ administration.

### 6.4. Neural atrophy mechanisms of A-T, in the context of other cerebellar ataxias

A-T is the most frequent variant in a series of autosomal recessive cerebellar ataxias, where a dysfunctional disease protein results in impaired DNA repair and a neural atrophy ensues, sometimes also accompanied by elevated blood levels of the fetal osmotic regulator AFP (Renaud et al., 2020; Synofzik et al., 2019). Thus, it is conceivable that the cascade of molecular events elucidated in A-T has a relevant overlap with the pathogenesis mechanisms of these ataxias.

Regarding the unrepaired DNA damage of A-T patients, it has been observed that retrotransposons cause DSB, that ATM modulates retrotransposon activation, and that this phenomenon in cerebella is a frequent cause of the manifestation of ataxia (Coufal et al., 2011; Gasior et al., 2006; Takahashi et al., 2022; Zhao et al., 2023). It is thought that genome modification by retrotransposons and its repair by ATM had a crucial role in animal phylogenesis when mammals developed placentas, modifying a retrotransposon sequence to serve in the regulation of the fetal capillary network and the endocrine signaling needed for eutherian intrauterine development (Kaneko-Ishino and Ishino, 2023; Rawn and Cross, 2008). It is therefore very interesting that the RTL1 or PEG11 protein, as one of these retrotransposon-like gene products, is among the strongest downregulations in the ATM-kd neuroblastoma cells, both in the ATM/ATR survey and in the global proteome profile. RTL1 also modulates the axon pathfinding needed for the development of brain hemisphere commissures (Kitazawa et al., 2021), so it represents a potential molecular bridge between the unrepaired DSBs, dysregulation of capillary blood vessels, and impaired axonal pruning that are consequences of ATM dysfunction.

It would be interesting to assess if decreased secretion of CHGA into the extracellular fluids is also observed in other cerebellar ataxias with elevated AFP blood levels. In proteome profiles of CSF from Parkinson’s disease (PD) patients, both chromogranins and secretogranins were found decreased, suggesting that the vesicle dynamics impairment well known to underlie PD includes a deficient secretion of secretory granules and/or synaptic LDCVs (Rotunno et al., 2020). In contrast, the immunohistochemical analysis of motor neurons in the spinal cord from patients with Amyotrophic Lateral Sclerosis revealed an accumulation and aggregation of CHGA (Schrott-Fischer et al., 2009).

Finally, some autosomal recessive cerebellar ataxias are characterized by dyslipidemia (Synofzik et al., 2019). Our observations of high APOB, LDLR and RELN abundance in the ATM-kd neuroblastoma cells, with previous observation of elevated APOB but low RELN in A-T patient CSF (Canet-Pons et al., 2018), is reminiscent of other childhood cerebellar ataxias that are caused by the loss of APOB or mutant RELN-VLDLR (Hentati et al., 2012; Homer et al., 2005; Valence et al., 2016). Overall, the depletion of ATM affects DNA repair, secretion of proteins like AFP, and lipid homeostasis as pathways that play well-established roles in the pathogenesis of ataxias.

## 7. Conclusion

In this study, we describe novel functions of cytoplasmic ATM in the regulation of vesicular secretion via CHGA, and of axonal pruning via semaphorin/ephrin-CRMP5-SPAST signaling. Phenotypically, ATM-kd cells exhibited significantly retracted neurites with excessive microtubule stabilization **(Fig. 8)**. Our observations suggest additional molecular biomarkers for the clinical assessment of A-T patients and may become useful in understanding the molecular pathogenesis also in other variants of autosomal recessive cerebellar ataxias.

**Figure 8:**
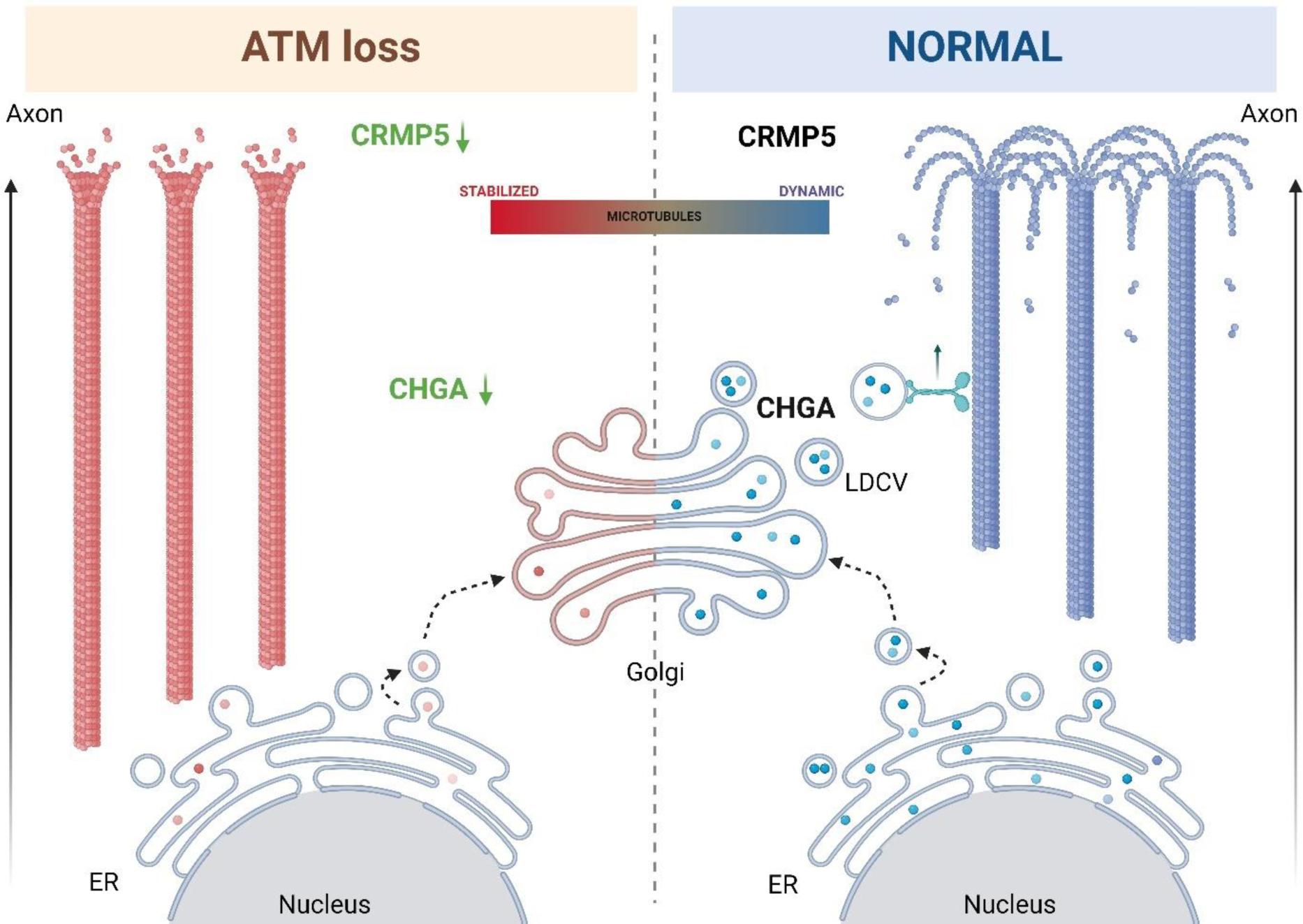
Proposed pathway for the novel cytoplasmic ATM-mediated signaling events. Compared to normal cells with neural origin (right panel), CHGA protein levels and secretion is reduced and CRMP5 protein levels as well as phosphorylation at Ser538 is diminished upon ATM loss (left panel). Given the crucial role of CHGA in dense-core granule biogenesis (Koshimizu et al., 2010), we propose that reduced CHGA in granules in the trans-Golgi network (TGN) diminishes budding of large dense core vesicles (LDCV) from TGN membrane resulting in reduced transport of LDCVs along the axon and reduced secretion of LDCVs to the extracellular space. This probably results in reduced neuropeptide and neurotrophic signaling. On the other hand, the concomitant CRMP5 protein loss causes increased polymerization of microtubules, probably via reduced binding and sequestration of tubulin heterodimers, shifting the microtubule equilibrium from dynamic towards stabilized, thereby causing neurite retraction. Abbreviations: ER = endoplasmic reticulum, LDCV = large dense-core vesicle, CHGA = chromogranin A, CRMP5 = Collapsin Response Mediator Protein 5.

## 10. Data availability

Data will be made available on request.

## 11. Author contributions

Conceptualization, M.R. and G.A.; Methodology, M.R., R.P.D., H.R., R.S., K.A., A.P.P., M.P.S. and G.A.; Software, J.K.; Validation, M.R. and G.A.; Formal analysis, M.R., J.K. and G.A.; Investigation, M.R., H.R., K.A., A.P.P., M.P.S. and G.A.; Resources, M.R., R.S. and G.A.; Data Curation, K.A., A.P.P. and M.P.S.; Writing - Original Draft, M.R. and G.A.; Writing - Review & Editing, M.R., R.P.D., H.R., J.K., R.S., K.A., A.P.P., M.P.S. and G.A.; Visualization, M.R., J.K. and G.A.; Supervision, R.S., M.P.S. and G.A.; Project administration, M.R. and G.A.; Funding acquisition, R.S. and G.A.

## 12. Acknowledgements

We thank Suzana Gispert, Stefan Momma and Tanja Müller for advice. We are grateful to the staff of the animal facility ZFE at the Goethe University Hospital for technical assistance. Sketches were created with BioRender.com.

## 13. Appendix. Supplementary data

Supplementary data to this article can be found online.

**Figure S1:**
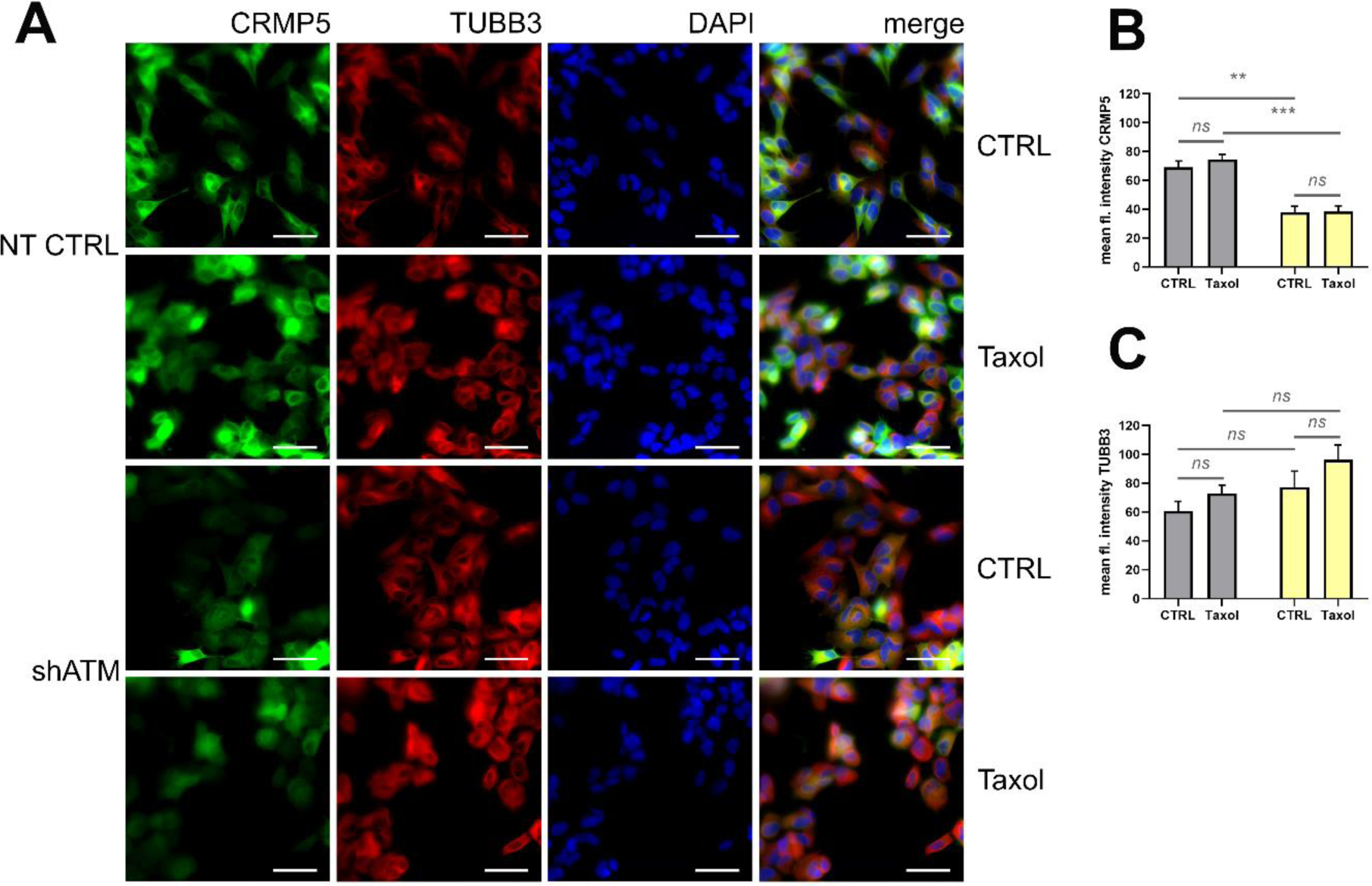
Immunofluorescence analysis of CRMP5 *in vitro.* **(A)** Immunofluorescence of CRMP5 (green), TUBB3 (red) and DAPI in unstressed or Taxol-treated SH-SY5Y cells. Images were taken at 60x magnification. Scale bar corresponds to 50 µm. **(B, C)** Quantification of fluorescence intensity using ImageJ after staining for **(B)** CRMP5 or **(C)** TUBB3. Fluorescence intensity was measured in three biological replicates each. Bar plots represent mean + SEM. Asterisks reflect significance: * = p ≤ 0.05, ** = p ≤ 0.01, *** = p ≤ 0.001, *** = p ≤ 0.0001; T = p ≤ 0.1, ns = non-significant.

**Figure S2:**
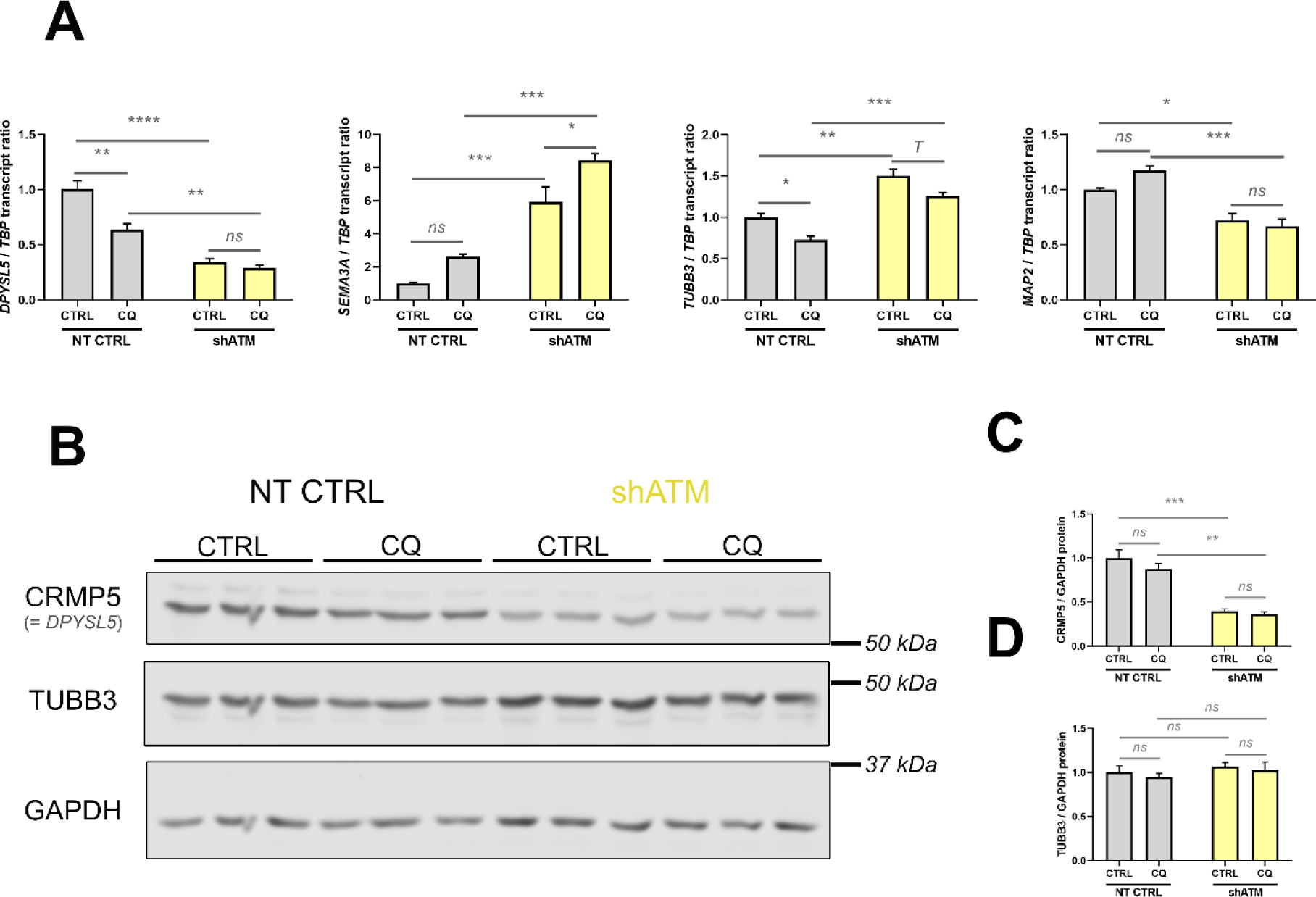
Dysregulation of the semaphorin-CRMP5 axis upon ATM loss and osmotic stress *in vitro.* **(A)** Significant reduction of *DPYSL5* (CRMP5 protein) transcript levels in shATM cells and during the *in vitro* osmotic stressor treatment with CQ. Significant induction of transcript levels of the upstream signaling receptor *SEMA3A*. Transcript levels of CRMP5 downstream signaling components *TUBB3* and *MAP2*. Transcript levels are n=3. **(B)** Immunoblots for quantification of the protein levels of ATM, CRMP5 and TUBB3 with GAPDH as housekeeping protein in an osmotic stress experiment using CQ. **(C)** Densitometric quantification of CRMP5 protein to GAPDH levels, with significant reduction in shATM cells. **(D)** Densitometric quantification of TUBB3 protein to GAPDH levels. Protein levels are n=3. Bar plots represent mean + SEM. Asterisks reflect significance: * = p ≤ 0.05, ** = p ≤ 0.01, *** = p ≤ 0.001, *** = p ≤ 0.0001; T = p ≤ 0.1, ns = non-significant. Abbreviations: CQ = chloroquine.

**Figure S3:**
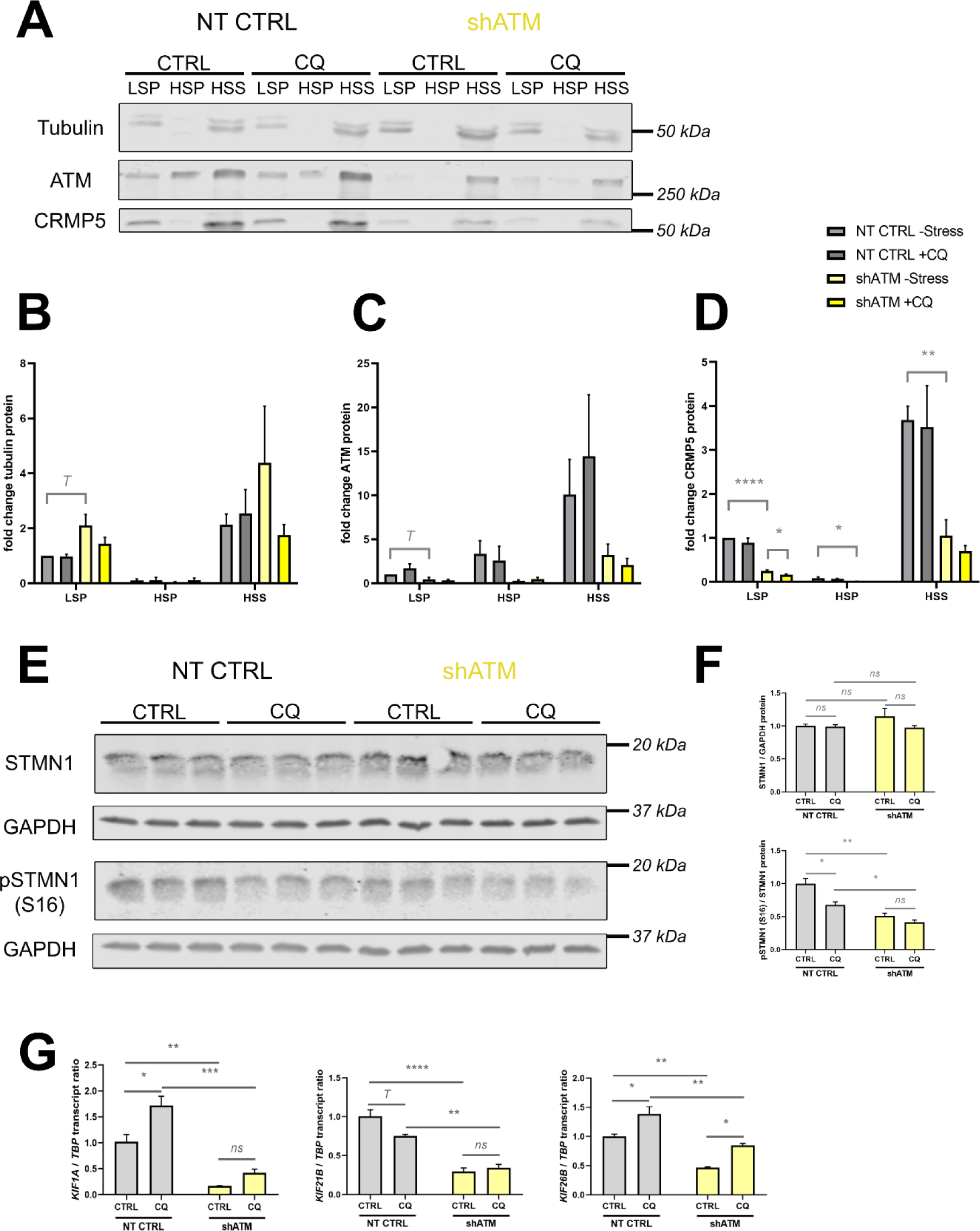
Osmotic stress does not alter microtubule dynamics *in vitro* but influences mRNA expression of microtubule-based anterograde axonal transport proteins. **(A)** Immunoblots for quantification of the microtubule content versus free tubulin and for quantification of ATM and CRMP5 protein in different fractions. **(B)** Densitometric quantification of tubulin protein in the different fractions. Tubulin protein shifted from the HSS fraction containing free tubulin towards the LSP fraction containing microtubules upon ATM-kd (as statistical trend), but not upon treatment with CQ. Please note that the unstressed samples correspond to the ones shown in Figure 5. **(C)** Densitometric quantification of ATM protein in the different fractions. ATM was detected mostly in the HSS fraction and was generally reduced in shATM cells. **(D)** Densitometric quantification of CRMP5 protein in the different fractions, following a similar pattern as ATM protein. Graphs are depicted as fold changes to the unstressed NT CTRL LSP. Protein levels are n=3. **(E)** Immunoblots for quantification of STMN1 and its phosphorylation at Ser16 as a second measure for microtubule stabilization (Jourdain et al., 1997). **(F)** Densitometric quantification of STMN1 total protein levels to GAPDH housekeeping protein and densitometric quantification of p-Ser16-STMN1 levels to STMN1 total protein, revealing decreased phosphorylation of STMN1 in shATM cells. Protein levels are n=3. **(G)** Transcript levels of different microtubule anterograde axonal transport factors *KIF1A*, *KIF21B* and *KIF26B*, in an osmotic stress experiment using CQ treatment. Transcript levels are n=3. Bar plots represent mean + SEM. Asterisks reflect significance: * = p ≤ 0.05, ** = p ≤ 0.01, *** = p ≤ 0.001, *** = p ≤ 0.0001; T = p ≤ 0.1, ns = non-significant. Abbreviations: LSP = low-speed pellet, HSP = high-speed pellet, HSS = high-speed supernatant, CQ = chloroquine.

